# PIPI-C: A combinatorial optimization framework for identifying post-translational modification crosstalks in mass spectrometry data

**DOI:** 10.1101/2024.08.06.606765

**Authors:** Shengzhi Lai, Shuaijian Dai, Peize Zhao, Chen Zhou, Ning Li, Weichuan Yu

**Affiliations:** Department of Electronic and Computer Engineering, The Hong Kong University of Science and Technology, Hong Kong, China; Department of Biomedical Engineering, School of Basic Medical Sciences, Central South University, Hunan 410013, China; Interdisciplinary Programs Office, The Hong Kong University of Science and Technology, Hong Kong, China; Ningbo No.2 Hospital, China, Zhejiang 315010, China

**Keywords:** Post-translational modification (PTM), PTM crosstalk, Peptide identification, Computational proteomics, Mixed integer linear programming (MILP)

## Abstract

Post-translational modifications (PTMs) are pivotal in cellular regulations, and their crosstalk is related to various diseases such as cancer. Given the prevalence of PTM crosstalk within close amino acid ranges, identifying peptides with multiple PTMs is essential. However, this task is an NP-hard combinatorial problem with exponential complexity, posing significant challenges for existing analysis methods. Here, we introduce PIPI-C (**P**TM-Invariant **P**eptide **I**dentification with a **C**ombinatorial model), a novel search engine that addresses this challenge through a mixed-integer linear programming (MILP) model, thereby overcoming the limitations of existing approaches that struggle with high-order PTM combinations. Rigorous validation across diverse datasets confirms PIPI-C’s superior performance in detecting PTM crosstalks. When applied to over 72 million mass spectra of three human cancers—lung squamous cell carcinoma (LSCC), colorectal adenocarcinoma (COAD), and glioblastoma (GBM)—PIPI-C reveals significantly upregulated PTM crosstalks. In LSCC, 50% of 860 upregulated unique PTM site patterns (UPSPs) (when comparing cancer vs. normal samples) carried at least two PTMs, including literature-supported crosstalks such as di-methylation with trifluoroleucine substitution and amidation with proline-to-valine substitution. Similar findings in COAD and GBM highlight PIPI-C’s utility in uncovering cancer-relevant PTM crosstalk landscapes. Overall, PIPI-C provides a robust mathematical framework for decoding complex PTM patterns, advancing our understanding of PTM-driven cellular processes in diseases.

## 1 Introduction

Post-translational modifications (PTMs) are crucial for regulating diverse cellular processes ^1^, including signal transduction, gene expression regulation, protein activity, and cell cycle control. To date, PTMs have been extensively observed across various organisms, underscoring their universal importance. The Eukaryotic Phosphorylation Site Database (EPSD) ^2^ has recorded 1,616,804 ex-perimental phosphorylation sites in 209,326 phosphoproteins across 68 eukaryotes, including Homo sapiens, Mus musculus, Drosophila melanogaster, etc. dbPTM^3^ has recorded over 2,790,000 PTM sites across organisms, with over 2,243,000 experimentally validated. Over the past three decades, extensive research has focused on the mechanisms of single PTMs^4–6^. However, PTMs do not function in isolation. Recently, increased attention has been directed toward understanding the combinatorial effects of multiple PTMs on proteins, i.e., a PTM can either promote or prevent the function of another PTM, collectively referred to as PTM crosstalk ^7–10^. Studies have shown that PTM crosstalk is associated with diverse human diseases, such as tumorigenesis^11^, Alzheimer’s disease ^12^, Parkinson’s disease ^13^, and cardiovascular disorders ^14^. For example, when analyzing 1,251 sirtuin interactors, Aggarwal et al. ^15^ found that approximately 83% of the proteins contain at least one competitive PTM crosstalk site, where different modifications compete for the same residue. Notably, half of these crosstalk sites were located in proteins associated with cardiovascular diseases (CVDs), highlighting their potential clinical relevance. Further, Wu et al. ^16^ summarized 25 typical examples of PTM crosstalks related to various types of cancers, including nasopharyngeal carcinoma, hematopoietic cancer, prostate cancer, colorectal cancer, breast cancer, etc. Despite these findings, research on PTM crosstalk has only revealed the tip of the iceberg. For example, PTMcode ^17^ predicted more than 1,450,000 potential PTM crosstalks based on evidence for residue competition, close structural distance, and coevolution. However, it has only recorded about 200 experimentally validated PTM crosstalks so far. Another database, dbPTM^3^, has also recorded only 491 experimentally validated PTM crosstalks.

PTM crosstalk can occur inter- or intra-protein^18,19^ when PTMs are structurally close in space due to protein-protein interactions or protein folding. More importantly, multiple PTMs can occur on adjacent amino acid residues in the primary sequence, termed “PTM hot-spot” ^20,21^, which facilitate intra-protein PTM crosstalk within these regions.

Crosstalk within PTM hot-spots is commonly observed. Aggarwal et al. ^15^ identified 614 proteins containing PTM hot-spots and 133 with crosstalk hot-spots by integrating multiple databases in a cardiovascular disease study. A well-known example of hot-spot PTM crosstalk is the “histone code” ^22^, i.e., patterns of co-existing modifications on histones, such as acetylation, methylation, phosphorylation, ubiquitination, and malonylation ^23^. Histone proteins and peptides are extensively modified: Taylor et al. ^24^ showed that in untreated SUM159 cell data^25^, 95% of H4 proteins carried at least two PTMs, with 45% bearing three to five. From a mouse stem cell dataset, Schwämmle et al. ^26^ identified numerous multiply modified histone peptides, with 70% carrying two to five PTMs. Another example is the “CTD (C-terminal domain) code” ^27,28^ on RNA polymerase II, where the conserved heptapeptide repeat YSPTSPS undergoes multiple phosphorylations or glycosylations during gene regulation. In practical applications, hot-spot PTM crosstalk links to human diseases: García-Moreno et al. ^29^ showed that the peptide CFFCHAP, bearing three RA-associated PTMs, acts as a rheumatoid arthritis (RA) antigen. Notably, severe joint damage correlated significantly with IgG isotype only when the tri-modified peptide was used. However, the peptide and PTMs were selected based on prior RA knowledge, a limitation that hinders generalization to other diseases.

Current PTM crosstalk research relies heavily on datasets enriched for multiple targeted PTMs^16,30–32^. Researchers acquire such data using techniques like simultaneous ^30,33^ or serial ^34^ enrichment of multiple PTMs, though the selection of the target PTMs remains limited relative to the over 200 known PTM types ^35^. Alternatively, researchers leverage existing PTM datasets ^17,26^ to identify crosstalk, a process requiring access to large-scale resources.

In proteomics research, diverse protein identification techniques have been developed since the 1950s, including Edman degradation, recognition tunneling, nanopore technology, and mass spectrometry (MS). Among these, MS is the most widely applied technique for high-throughput protein identification^36,37^. In MS-based bottom-up proteomics, protein samples are first digested into peptides before injection into the instrument, generating hundreds of thousands of mass spectra. Ideally, each spectrum corresponds to an unmodified, singly modified, or multiply modified peptide. Combined with the above discussion on PTM crosstalk in PTM hot-spots, identifying multiply modified peptides from these mass spectra is critical for enabling proteome-scale PTM crosstalk research. However, this remains challenging using existing database search methods^38,39^. More importantly, these PTMs must be identified simultaneously on the same protein molecule (rather than aggregating results from separate analyses), as proteins can be modified at distinct stages of their lifecycle (temporal dynamics) or in different subcellular locations, tissues, or organs (spatial distribution) ^40,41^. Localizing PTMs at adjacent sites in the same protein sequence captures only spatial information, which does not guarantee crosstalk; identifying multiple PTMs on the same peptide, however, captures both spatial and temporal context—a prerequisite for validating PTM crosstalk.

Identifying peptides with multiple PTMs can be framed as a combinatorial problem: determining the optimal PTM combination in the peptide backbone sequence, with which the theoretical MS2 spectrum is most similar to the experimental MS2 spectrum ^42^. For a peptide sequence consisting of *N*_*a*_ amino acids, the number of all possible *k*-PTM combinations, *N*_*c*_, can be calculated as

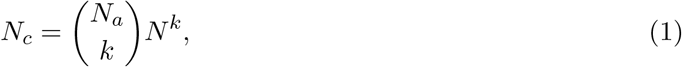

where *N* represents the average number of potential PTMs on each amino acid (as documented in the Unimod database). *N*_*c*_ increases exponentially with *k*, assuming at most one PTM on one amino acid. For instance, given *N*_*a*_ = 20, *k* = 4 and *N* = 100, this number reaches the order of 10^11^. Moreover, characterizing PTMs requires matching each MS2 spectrum to all peptide candidates in the database, a computationally intensive task due to the exponential increase in possible combinations.

Combinatorial problems are NP-hard due to their exponential complexity. They are typically solved using algorithms that involve a trade-off between computational complexity and solution optimality, such as exhaustive enumeration, heuristic algorithms, greedy algorithms, and exact algorithms. Since the 1990s, researchers have been developing such algorithms to identify peptides with multiple PTMs.

Initially, conventional database search methods (referred to as closed search), such as Mascot ^43^ and Comet ^44^, were among the first to identify peptides with a limited number (typically, nine) of pre-specified PTMs. These methods are essentially exhaustive methods within a limited scope, as the number of allowed predefined PTMs is highly insufficient relative to the 1,559 PTM entries documented in Unimod ^45^ (up to 23 Jan 2025). A rough estimation shows that the total number of all nine-PTM combinations reaches the order of 10^23^, not to mention that many PTMs can have multiple sites, which should be pre-specified as distinct PTMs.

Since 2010, open search methods have been developed for this combinatorial problem of identifying peptides with multiple PTMs. Most of these methods rely on short, PTM-invariant amino acid sequences (termed “tags”), which are unaffected by shifted precursor masses. By leveraging the difference between the theoretical mass of a peptide candidate and the precursor mass of an MS2 spectrum, these methods use alignment and enumeration strategies to characterize PTM patterns—specifically, the number, mass, and sites of PTMs on the peptides.

Alignment-based methods typically use dynamic programming. The MODplus series methods^46,47^ and PIPI^48^ employ dynamic programming to align extracted tags to peptide sequences and characterize PTMs. TagGraph ^49^ uses dynamic programming to align PTM-containing peptide candidates with *de novo*–generated sequences and localize multiple PTMs. Enumeration-based methods exhaustively analyze different PTMs or MS2 peaks. PeaksPTM ^50^ exhaustively enumerates peptides with common or unspecified PTMs and combines results to detect multiple PTMs. Open-pFind ^51^ characterizes PTMs by enumerating all single PTMs or two-PTM combinations. MSFragger^52,53^ enumerates all possible modified locations using peaks matching regular or shifted ions to localize PTMs. In summary, alignment-based methods typically identify one PTM within a gap between two tags, and their performance heavily relies on the correctness of the tags. Enumeration-based methods often limit their scope to peptides with single PTMs or two-PTM combinations, as handling higher numbers becomes computationally intractable. These limitations have underscored the need for a more theoretical understanding of this combinatorial problem to find the optimal solution.

Integer linear programming (ILP) is a mathematical framework for solving combinatorial problems, offering guaranteed optimality. It has been utilized in computational proteomics ^54–56^. He et al. ^57^ constructed a partial set covering model for protein mixture identification. Mitsos et al. ^58^ and Ji et al. ^59,60^ employed ILP to infer cell-specific signaling pathway networks for drug efficacy prediction. DiMaggio et al. developed ILP models for de novo peptide identification^61^ and for the identification of targeted PTMs in known backbone sequences^62^. Later, Baliban et al. ^63^ combined and extended these two models to detect untargeted PTMs using de novo sequences derived from small-scale, low-accuracy MS2 spectra. In our previous work, PIPI2^64^, we employed a greedy approach to solve this combinatorial problem, though it lacked optimality guarantees. Here, we formulate a mixed integer linear programming model (MILP) to achieve the optimal solution. MILP can be solved using various methods ^65–68^, which are implemented in high-performance solvers such as Gurobi^69^.

In this paper, we propose PIPI-C (**P**TM-**I**nvariant **P**eptide **I**dentification with a **C**ombinatorial model), a search engine that utilizes a novel MILP model to identify peptides with multiple PTMs. PIPI-C uses a fuzzy and bidirectional matching approach to locating tags in protein sequences. It then constructs and solves the MILP model to determine the peptide sequences and the PTM patterns. To the best of our knowledge, PIPI-C provides the first combinatorial optimization model for identifying peptides with multiple PTMs. First, we comprehensively validated PIPI-C’s performance using various types of data sets and compared it to two existing software programs. Then, we applied PIPI-C to over 72 million mass spectra of three different human cancers: lung squamous cell carcinoma (LSCC), colorectal adenocarcinoma (COAD), and glioblastoma (GBM) to identify significantly upregulated PTM crosstalks in these cancers.

## 2 Results

### 2.1 Performance validation

To evaluate the performance of PIPI-C on backbone (peptide sequence, disregarding PTMs) identification and PTM characterization, we conducted four experiments, comparing it with Open-pFind ^51^ (version 3.2.0) and MODplus ^47^ (version 2.01), which are recognized as leading tools for identifying peptides with PTMs. As shown previously ^64^, other programs, such as MSFragger ^52^, PeaksPTM ^50^, and TagGraph ^49^ exhibit limitations in their ability to handle peptides with multiple PTMs. Most parameters were set to default values except for specific ones outlined in **Supplementary Ta-ble S3**. Original parameter files, results, and simulation data sets are deposited on Zenodo at https://doi.org/10.5281/zenodo.15744650. All other data sets are available online (see **Data Availability**).

### 2.2 Performance validation with simulated data sets

In the first experiment, we evaluated the precision and sensitivity of backbone identification and PTM characterization of PIPI-C using seven simulated data sets with different signal-to-noise ratios (SNRs) generated by AlphaPeptDeep ^70^. The ground truth of the peptide backbone and PTM pattern of every MS2 spectrum is known. In each MS2 spectrum, the intensities of *b* and *y* ions were predicted using the protein Packet Kmod as a template (from ProteomeTools^71^ on ProteomeXchange Consortium^72^ with the data identifier PXD009449). We randomly appended noise peaks to the predicted signal peaks to create full MS2 spectra and finally obtained seven data sets with different SNRs, as detailed in **Supplementary Note 1**. Each data set consisted of 124,248 MS2 spectra with up to four PTMs from 18 different selected PTMs. These MS2 spectra were distributed as follows: 135 0-PTM spectra, 20,004 1-PTM spectra, 92,357 2-PTM spectra, 11,263 3-PTM spectra, and 489 4-PTM spectra.

In the evaluation, a correct backbone identification refers to a PSM where the backbone sequence is correct regardless of PTMs. Within the correct backbone identifications, a correct PTM characterization denotes a PSM with both the accurate backbone and the correct numbers, types, and sites of PTMs. For quality control, the false discovery proportion (FDP) was directly calculated based on the known ground truth. However, since the target-decoy strategy^73^ is mandatory in Open-pFind, we still added decoy proteins as extra irrelevant proteins in the database. The results were filtered at a peptide-level FDP of 0.01 for all search engines.

In the data set with SNR = 2.023, PIPI-C identified over 99% of all MS2 spectra with correct peptide backbones, while the values for Open-pFind and MODplus were 42% and 92%, respectively (**Fig. 1A**). Among these identifications, PIPI-C correctly characterized two times and 21% more PTM patterns than OpenpFind and MODplus, respectively. Notably, PIPI-C exhibited the highest PTM characterization precision, 85%, while the values for Open-pFind and MODplus were 65% and 76%, respectively. As the SNR decreased, both PIPI-C and Open-pFind maintained relatively consistent performances, whereas MODplus’s performance degraded more quickly (**Fig. 1A**). The PSMs with correct PTM patterns were categorized into four groups based on the true numbers of PTMs in the peptides (**Fig. 1B**). PIPI-C demonstrated superiority over competitors, especially for peptides with more than two PTMs. Open-pFind’s performance declined when peptides had more than one PTM, with near-zero correct identifications for peptides containing three or four PTMs. MODplus also failed to characterize 4-PTM peptides. Regarding each simulated PTM, both PIPI-C and MODplus displayed more uniform sensitivity (the ratio of the identified number over the truth number of a PTM) than Open-pFind, as illustrated in **Fig. 1C**.

**Figure 1:**
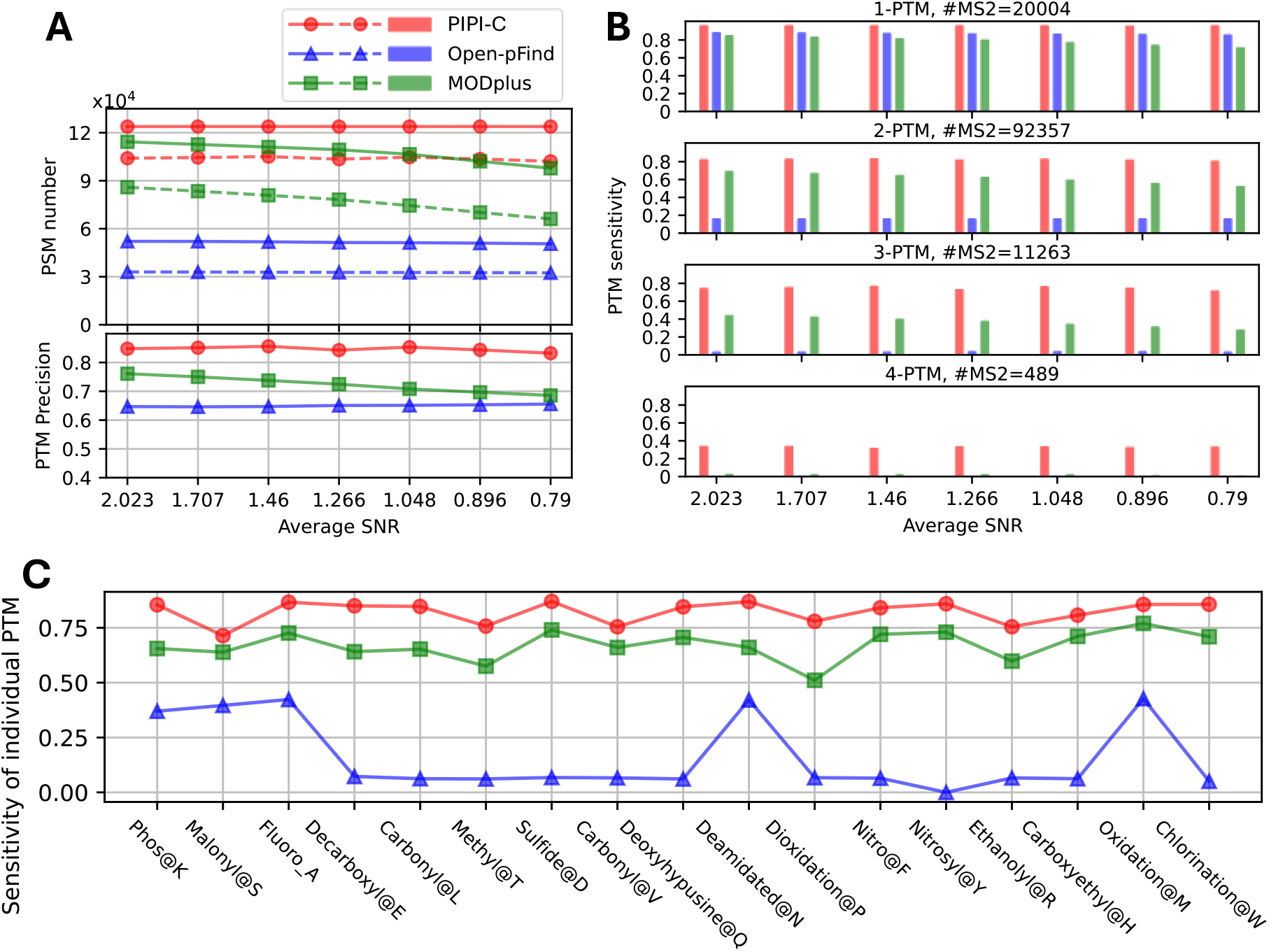
Comparison results using simulated data sets. PIPI-C performed better than competitors, especially for peptides with more than two PTMs. (A) Upper subplot, the number of PSMs with correct backbones (solid lines) and the number of PSMs with correct PTM patterns (dashed lines); lower subplot, the precision of PTM characterization, calculated by the number of PSMs with correct PTM patterns divided by the number of PSMs identified as carrying PTMs. (B) Sensitivity of PTM characterization, calculated by the number of PSMs with correct PTM patterns divided by the ground truth number of MS2 spectra with PTMs. (C) PTM characterization sensitivity for individual PTMs.

In our previous work ^64^, we observed that Open-pFind and MODplus incorporate “tolerances” for two user-defined parameters, namely enzyme specificity and maximum missed cleavages, which may compromise their performance in backbone identification when applied to the simulated data sets. We removed the influenced PSMs; otherwise, the performance would have been worse.

We also performed a negative control test using the simulated data sets and an irrelevant protein database. In this case, we search the data set with SNR = 2.023 against an *E*.*coli* protein database, which is irrelevant to the template proteins. The minimum q-value of the results is 0.33, which means PIPI-C has no identifications. This indicates that PIPI-C maintains a high level of specificity, and the false positives are well controlled when presented with unrelated protein sequences.

#### 2.1.2 Performance validation using synthetic data sets

Since simulation-based validation heavily depends on the simulation model and sometimes may not be realistic, we also use more realistic datasets for evaluation. In the second experiment, we evaluated PIPI-C using 21 synthetic data sets ^71^. The data sets contain 21 raw files of 1,023,540 MS2 spectra in total. Within each raw file, the MS2 spectra were generated from 115 to 200 unique peptides with at most one selected PTM on the selected amino acid, as detailed in **Supplementary Table S4**. Based on the target-decoy strategy, the results were filtered at a peptide-level false discovery rate (FDR) of 0.01 calculated by (*N*_*d*_ + 1)*/N*_*t*_ ^74^, i.e., the number of decoy matches plus one over the number of target matches.

We exemplify the result with the data set containing Phospho@Y (phosphorylation on Y, similar for the following abbreviations) in **Fig. 2** and record all 21 results in **Supplementary Fig. S1, Fig. S2**, and **Fig. S3**. The majority of identified backbones are from the selected template protein. At the same time, the second and third largest fractions come from irrelevant proteins for quality control (QC) and retention time (RT) calibration. Compared to Open-pFind and MODplus, PIPI-C demonstrated higher backbone sensitivity by respectively identifying 7% and 15% more PSMs and higher PTM sensitivity by characterizing more Phospho@Y. Open-pFind characterizes PTMs by enumerating single PTMs and 2-PTM combinations, which is advantageous in data sets featuring at most one PTM. Nonetheless, PIPI-C is still better than Open-pFind on these data sets.

**Figure 2:**
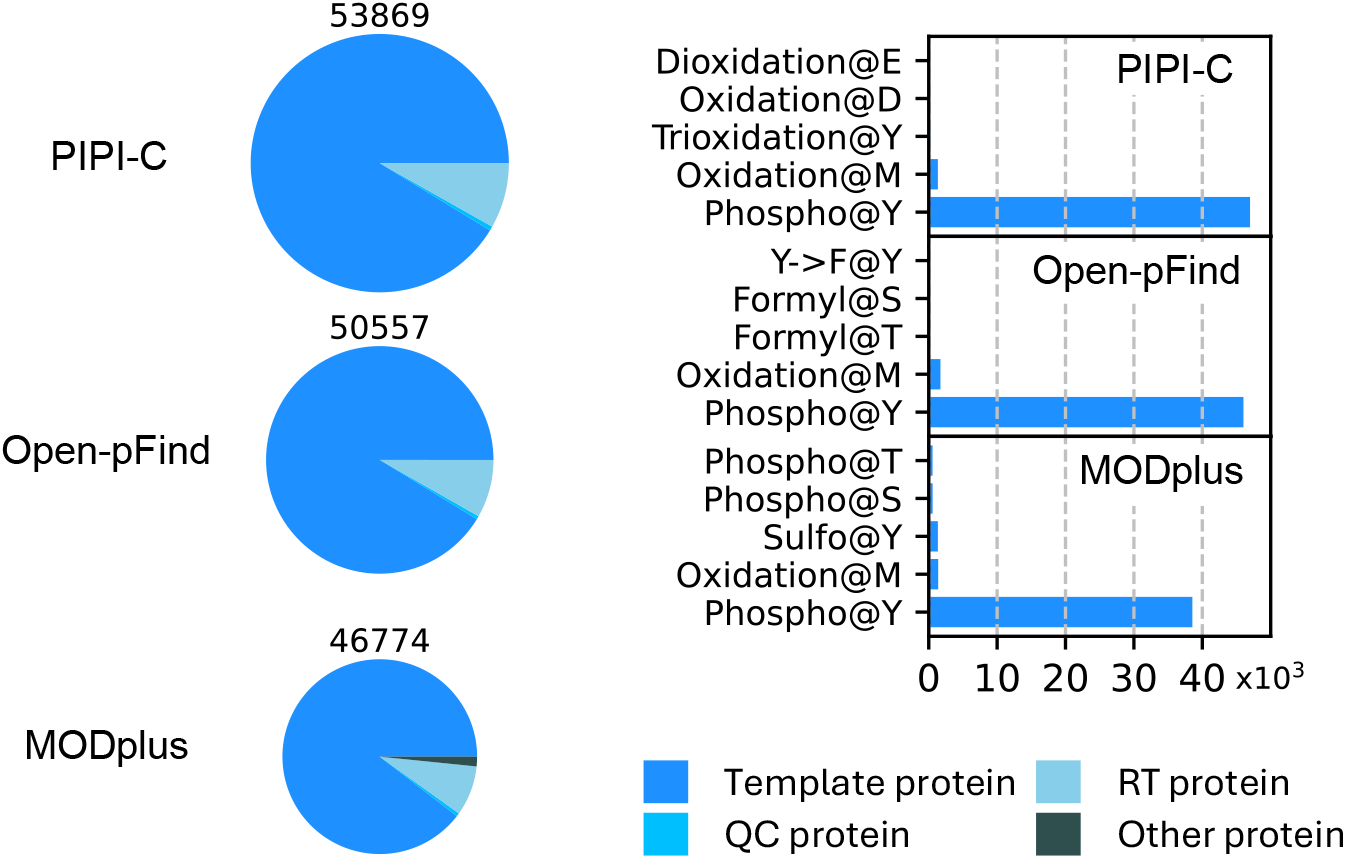
Comparison result of the synthetic data set with Phospho@Y at FDR of 0.01: bar chart, PSM numbers; pie chart, PTM numbers. On this data set with low complexity (up to one selected PTM), PIPI-C still demonstrated better performance than competitors. QC (quality control) and RT (retention time calibration) proteins are from the original data and are irrelevant to our study.

#### 2.1.3 Performance validation using dimethyl-labeled soybean data sets

In the third experiment, we evaluated PIPI-C using soybean data sets ^75^. These data sets consisted of two groups of raw files, collectively comprising approximately 800,000 MS2 spectra. Each group of raw files corresponded to fractionated peptides labeled with either light (L) dimethyl (monoisotopic mass 28.031 Da) or heavy (H) dimethyl (monoisotopic mass 34.063 Da). Both types of dimethyl labels could occur at any Lysine (K) or peptide N-term. No peptide contained both light and heavy dimethyl labels. We treated these two types of labels as unspecified PTMs without setting them as variable modifications and conducted searches against a soybean database that was provided alongside the data set.

The performance evaluation of PIPI-C, Open-pFind, and MODplus was based on the number of peptide-spectrum matches (PSMs) where the peptides were fully labeled by either light or heavy dimethyl labels. A peptide is considered fully labeled if both the N-term and all Ks in the sequence are labeled by either light or heavy dimethyl. These results were also filtered at a peptide-level FDR of 0.01, as in the second experiment. PIPI-C identified 75% and 17% more fully labeled peptides than Open-pFind and MODplus, respectively. Additionally, a significant intersection was observed among the three software programs, and PIPI-C uniquely identified a substantial number of peptides (**Fig. 3**).

**Figure 3:**
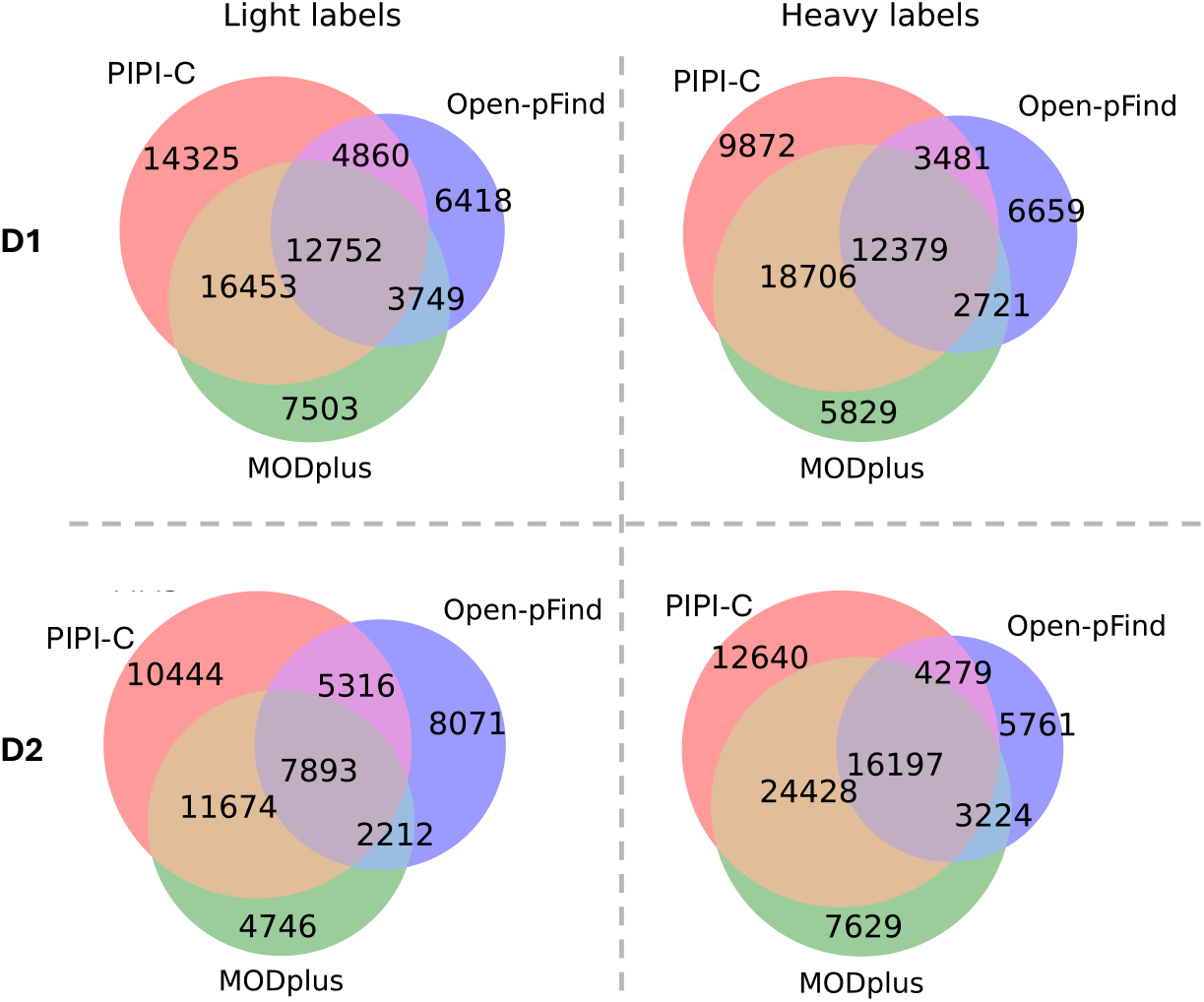
Intersection of fully dimethyl-labeled peptides identified from the soybean data sets R01. D1, D2: two samples in the data set.

During the data set preparation step, no peptide was labeled by both light and heavy dimethyl, which allowed us to estimate the error level of PIPI-C and MODplus using the ratio of the number of PSMs with peptides carrying both light and heavy labels over the number of PSMs involving the aforementioned fully labeled peptides. The ratios of PIPI-C and MODplus were 0.28% and 0.14%, respectively. Since all peptides identified by Open-pFind contain at most one dimethyl label (at the peptide N-term), the error level of Open-pFind could not be estimated in the same way. Results for the other five data replicates are shown in **Supplementary Fig. S4** to **Fig. S8**. PIPI-C consistently outperformed Open-pFind and MODplus in these replicates.

#### 2.1.4 Showcase of identifying PTM combinations using the *Petunia* data sets

In the fourth experiment, we showcased the power of PIPI-C in identifying peptides with multiple PTMs using *Petunia* data sets ^76^ with an enriched PTM, GG@K. Rather than considering it as a variable modification, we treated GG@K as unknown and used it as a reference point to identify reliable PTM combinations. The three fractionated raw files (CK1 LF1.raw to CK1 LF3.raw) from a complete sample were jointly searched against a *Petunia hybrida* database provided alongside the data set. Oxidation@M and Acetylation@AnyN-term were set as variable modifications, and GG@K was treated as unexpected. The results were filtered at a peptide-level FDR of 0.01.

PIPI-C identified a total of 14,850 PSMs, surpassing Open-pFind and MODplus by 17% and 12%, respectively. Among these PSMs, PIPI-C detected 9,462 GG@K, which was 10% and 25% more than Open-pFind and MODplus, respectively (**Fig. 4A**).

**Figure 4:**
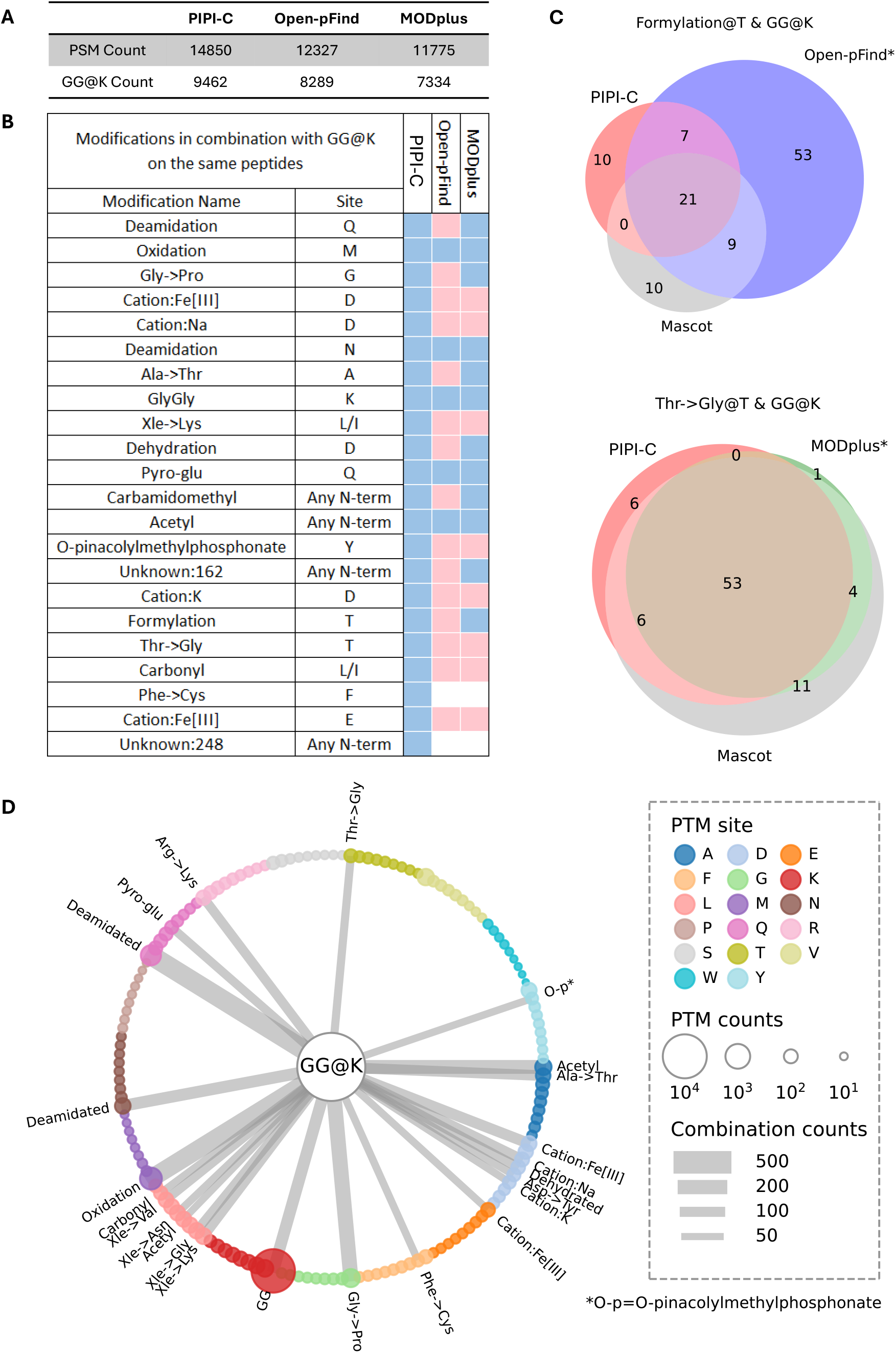
Results of the *Petunia* data sets. PIPI-C demonstrated better performance in identifying PTM combinations. (A) Number of PSMs and enriched GG@K identified using PIPI-C, Open-pFind, and MODplus. (B) PTMs co-occur with GG@K identified by PIPI-C, Open-pFind, and MODplus. Each PTM combination was identified more than 50 times by PIPI-C. (C) For Open-pFind and MODplus, unidentified PTM combinations can also be identified when the other PTM is pre-specified as a variable modification. (^*∗*^) denotes the search engine pre-specifying the PTM other than GG@K as a variable modification (D) PTM combinations with GG@K identified by PIPI-C more than 50 times. GG@K is visually placed in the center, connected to other PTMs in combination that are placed in the circle. The size of the dots indicates the count of the PTMs, and the strength of the connections indicates the count of the PTM combinations.

PIPI-C identified many PTM combinations involving GG@K and another PTM in the same peptides. Each of these PTM combinations was consistently identified in over 50 PSMs. As shown in **Fig. 4B**, some PTM combinations were consistently identified by the three search engines (blue cells), while others were not (red and white cells). For Open-pFind and MODplus, we manually set the other PTM in each unidentified PTM combination as a variable modification and searched again (one PTM each time). Most unidentified PTM combinations could then be identified (red cells). These results were further validated using Mascot ^43^ by setting both PTMs in each unidentified PTM combination as variable modifications (one combination at a time), as shown in **Fig. 4C** and **Supplementary Fig. 4C** where a single asterisk (^*∗*^) denotes the search engine pre-specifying the PTM other than GG@K as a variable modification. PIPI-C kept the same setting in all these comparisons without the variable modifications. While the extent of overlap varied among different PTM combinations, our analysis concludes that PIPI-C identified unspecified PTM combinations more effectively than Open-pFind and MODplus. **Fig. 4D** presents all the PTM combinations involving GG@K that have been consistently identified by PIPI-C in over 50 instances.

Moreover, PIPI-C successfully identified a 3-PTM combination of GG@K, Phe→Cys@F, and Unknown: 248@N-term (chemical composition: H(28) C(13) O(4)) in over 50 PSMs. In the column for Open-pFind and MODplus in **Fig. 4B**, two white cells indicate that by setting either PTM as a variable modification, Open-pFind and MODplus still could not detect the corresponding PTM combinations. However, when both PTMs were set as variable modifications in a single run, Open-pFind successfully identified the three-PTM combination, whereas MODplus did not. This was also verified by Mascot.

### 2.2 Application to LSCC data

#### 2.2.1 Results of LSCC1 data

We applied PIPI-C to a proteome-scale LSCC data set ^77^ generated by CPTAC, denoted as LSCC1. The data set consists of 22,121,363 tandem mass spectra (MS2 spectra) prospectively collected from 22 TMT-11 plex experiments from 108 treatment-naive, primary LSCC tumors, and 99 paired normal adjacent tissues, and labeled with TMT-11 for quantification analysis. The data search took about 12 days on a 32-core server.

At a peptide-level false discovery rate ^74^ (FDR) of 0.01, PIPI-C identified 10,616,210 peptide-spectrum matches (PSMs) with TMT-11 labels, among which over 50% carried PTM(s) besides the TMT-11 labels. Unique modified peptides followed Zipf’s law (**Fig. 5A**), and common PTMs such as Oxidation, Deamidation, and Formylation were abundantly identified (**Fig. 5B**). We cross-validated PIPI-C’s PTM search results using Mascot. We selected the top five major PTMs that have been widely studied ^3^, namely, phosphorylation, acetylation, GG, succinyl, and methylation, together with three less common PTMs, and two frequently identified amino acid substitutions, Xle to Pro and Xle to Asn. We collected the MS2 spectra from which these PTMs were identified, and searched them using Mascot by specifying each of these PTMs together with the TMT-11 label as variable modifications. The intersections between PIPI-C’s results and Mascot’s results are shown in **Fig. 5C**. It demonstrated that Mascot and PIPI-C have strong intersections on some of these PTMs, while some intersections are not significant. The reason we suppose could be the different scoring strategies of these two methods. Please note that Mascot is the state-of-the-art search engine with the best search performance if up to nine types of PTMs can be specified before database search. However, without pre-specification, Mascot cannot identify any of these PTMs. While it is challenging for Mascot to know which PTMs to search, our PIPI-C provides exactly such information. In this sense, the combination of PIPI-C followed by Mascot search is an enabling validation test design that combines the strengths of two different search engines. We further performed quantification analysis on the PSMs that were repetitively identified from at least 16 pools out of the total 22 pools. The quantification results (**Supplementary Table S5 and S6**) with q-value *<* 0.05 show that 4,593 unique PTM site patterns (UPSPs, groups of peptides containing the same PTM sites) have at least two PTMs, corresponding to 171,960 PSMs in total (**Table 1**). The PSMs with at least one PTM were subject to quantification analysis. Among all UPSPs, 860 were significantly upregulated (q-value *<* 0.05, fold change *>* 3), and 20 were significantly downregulated (q-value *<* 0.05, fold change *<* 1/3) in the lung cancer sample as compared to normal samples (**Fig. 6A**).

**Table 1:** UPSPs with different numbers of PTMs identified from LSCC1 data. 4593 UPSPs have at least two PTMs.

| PTM number<br>in UPSP | Total number<br>of UPSPs | Total number<br>PSMs | Average<br>pool number |
| --- | --- | --- | --- |
| 0 | 29326 | 2350669 | 19.2 |
| 1 | 9311 | 418681 | 17.5 |
| 2 | 4456 | 165632 | 16.8 |
| 3 | 108 | 4399 | 16.4 |
| 4 | 29 | 1929 | 16.5 |

**Figure 5:**
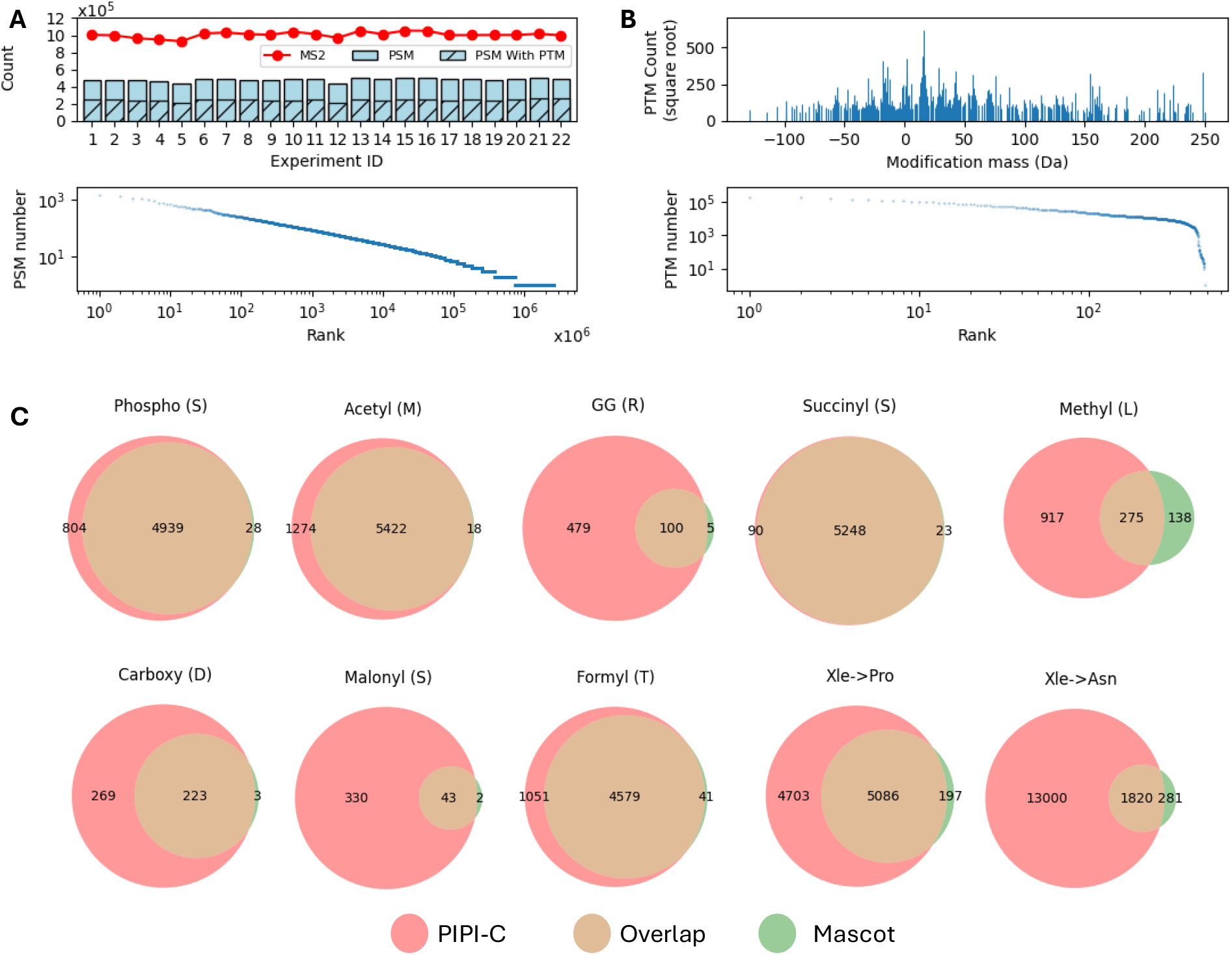
Search results of the LSCC1 data using PIPI-C at a peptide-level FDR *<* 0.01. (A) Identification results: upper, the number of PSMs and those with PTMs, and the number of MS2 spectra in each experiment; lower, PSMs with differently modified peptides ranked by the PSM number. (B) PTM mass distribution with some abundant ones annotated. (C) The overlap of results of PIPI-C and Mascot on several major PTMs and uncommon PTMs that are identified by PIPI-C.

**Figure 6:**
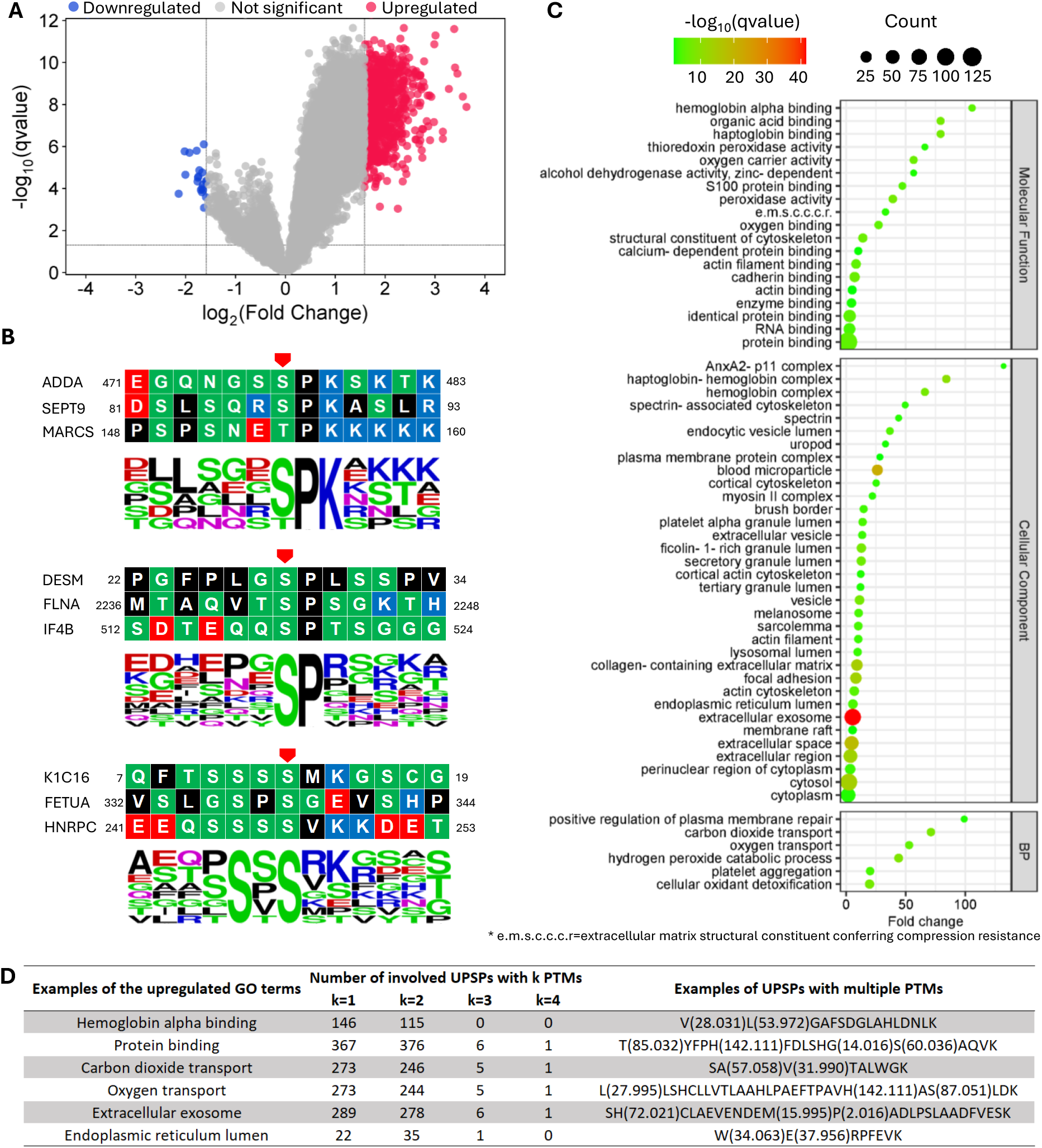
Quantification results and analysis of the LSCC1 data set, q-value *<* 0.05. (A) 860 upregulations (fold change *>* 3) and 20 downregulations (fold change *<* 1/3). (B) Sequence logos of upregulated Phosphorylated S/T, q-value *<* 0.01. (C) GO analysis of the proteins with significantly regulated UPSPs. BP: biological process, BH-FDR *<* 0.01. (D) Examples of the correspondence between the upregulated GO terms and UPSPs with different numbers of PTMs.

From the 860 upregulated UPSPs, we performed motif analysis and gene ontology (GO) analysis (**Supplementary Table S7**). We demonstrated several significantly enriched motifs, namely, p[S/T]PK, p[S/T]P, [S]xp[S/T] with q-value *<* 0.01 (**Fig. 6B**). Based on the GO analysis of the upregulated UPSPs, we found significant upregulations (q-value *<* 0.01) of biological process, cellu-lar component, and molecular function (**Fig. 6C**). Most of these upregulations are related to lung functions and the respiratory system and have been reported in the literature, such as hemoglobin alpha binding ^78^, protein binding ^79^, carbon dioxide transport ^80^, oxygen transport^81^, ANXA2^82^, extracellular vesicle ^83^, focal adhesion ^84^, and endoplasmic reticulum ^85^. Notably, the upregulation of extracellular exosomes had high statistical significance.

An example of the correspondence between UPSPs and the upregulated GO terms is shown in **Fig. 6D**. About 49% (420) of the involved UPSPs contain at least two PTMs, indicating extensive potential intra-peptide PTM crosstalks in LSCC1. These PTMs include methylation, ADPribosylation followed by conversion to hydroxamic acid via hydroxylamine, tyrosine oxidation to 2-aminotyrosine, etc. Among these PTMs, 63 are reported to be related to lung cancer, such as Benzyl isothiocyanate ^86^, quinone^87^, and monoglutamyl ^88^. The details of all the reference papers are deposited on Zenodo at https://doi.org/10.5281/zenodo.15744650.

Moreover, we analyzed the PPI network among the leading proteins of the 860 upregulated UPSPs (**Supplementary Table S8**). Applying the highest confidence (0.9) of required score and stringent FDR of 1%, we generated a full STRING network as shown in **Fig. 7**, where the LIFE (Likelihood of Interaction and Function Evaluation) score of each node is calculated as the summation of the log2 enrichment ratio, the number of GO terms corresponding to the protein, and the degree of the node in the figure. These interacting proteins correspond to GO terms such as antioxidant activity, oxygen carrier activity, blood coagulation, and blood microparticle (**Supplementary Table S9**). We identified two significant interacting groups, and most of the interacting proteins have been reported as lung cancer-related. For example, ZYX was found to be significantly elevated in the serum of NSCLC (non-small cell lung cancer) patients compared to healthy controls^89^, suggesting its potential as a diagnostic biomarker, although its levels did not correlate with tumor subtype, stage, or grade. In another study ^90^, Richter et al. identified ACTG1 as a recurrently amplified and overexpressed actomyosin gene across multiple cancer types, including lung cancer, suggesting its potential oncogenic role and relevance as a biomarker in lineage-specific cancer contexts. Furthermore, Katono et al. ^91^ showed that MYH9 is expressed in a subset of NSCLC tumors and is significantly associated with more aggressive features, such as poor differentiation, vascular and lymphatic invasion, and worse overall survival. Chen et al. ^92^ demonstrated that MYH9 enhances the stem cell-like properties of lung cancer cells by upregulating cancer stem cell markers (e.g., CD44, SOX2, Nanog) and activating the mTOR signaling pathway. Knockout or inhibition of MYH9 reduced these stemness traits, suggesting its role in maintaining tumor-initiating capacity. These examples show the power of PIPI-C in identifying cancer-related peptides.

**Figure 7:**
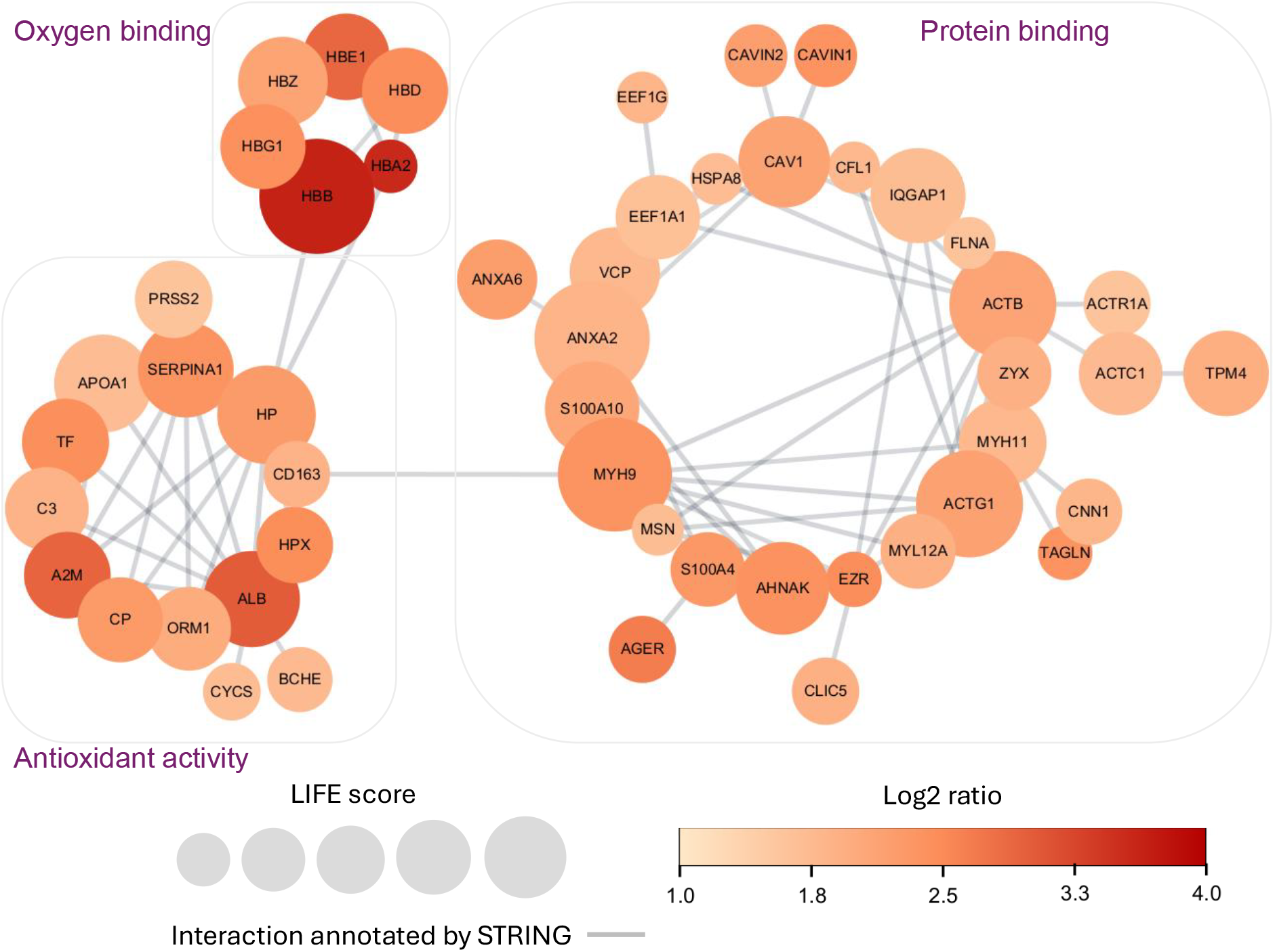
Interaction network among the proteins of the upregulated UPSPs. Two significant interacting groups were identified, with most of the interacting proteins being reported as lung cancer-related. Circular nodes represent significantly upregulated proteins. The size of the node represents the LIFE (Likelihood of Interaction and Function Evaluation) score calculated by log2-ratio, GO analysis results, and STRING interaction score. The edge represents a probable protein-protein interaction predicted by the STRING database. The thickness represents the confidence score (≥0.9) of interactions.

#### 2.2.2 Extra LSCC cohorts

To support the findings in the LSCC1 data, we used two extra LSCC cohorts (LSCC2, LSCC3) from other research groups as an independent validation. The first cohort ^93^, LSCC2, comprises 11,419,378 MS2 spectra collected from tumor tissues of 108 treatment-naive LSSC western patients. The data is organized into 29 batches (12 fractions in each batch) and labeled with TMT-6. The second cohort ^94^, LSCC3, comprises 3,147,669 MS2 spectra collected from tumor tissues of 25 treatment-naive Chinese patients with LSSC. The data is organized into five batches (20 fractions in each batch) and labeled with TMT-6. The search parameters of both cohorts are recorded in **Supplementary Table S3**.

At a peptide-level FDR of 0.01, we identified 1,333,749 with TMT labels from LSCC2, among which 560,334 carried PTM(s) other than the TMT labels; these two numbers for LSCC3 are 1,248,897 and 604,987. The modified peptides from LSCC1 can be validated by LSCC2 and LSCC3 data, as shown in **Fig. 8A**. The first column is the number of PTMs in the peptides identified in LSCC1. The third to sixth columns are the total number of these peptides, and those that are also identified in LSCC2, LSCC3, and both LSCC2 and LSCC3. The color bars in the cell represent their proportion in the total number of peptides (the third column). The results from LSCC1 can be supported by LSCC2 and LSCC3 data in two aspects. First, the rows of “UPSP” show that among UPSPs with one PTM, about 27% and 45% are identified in LSCC2 and LSCC3, respectively, and about 20% are identified in both. Second, since some UPSPs are identified in multiple PSMs, we show the proportion of PSMs that are also identified by LSCC2 and LSCC3, i.e., about 40% and 56% PSMs with one PTM identified in LSCC1 are identified in LSCC2 and LSCC3, respectively, and about 32% can be identified in both. While the proportions in the rows with three or four PTMs are lower, it still shows good overlap (about 20%) for the two-PTM row.

**Figure 8:**
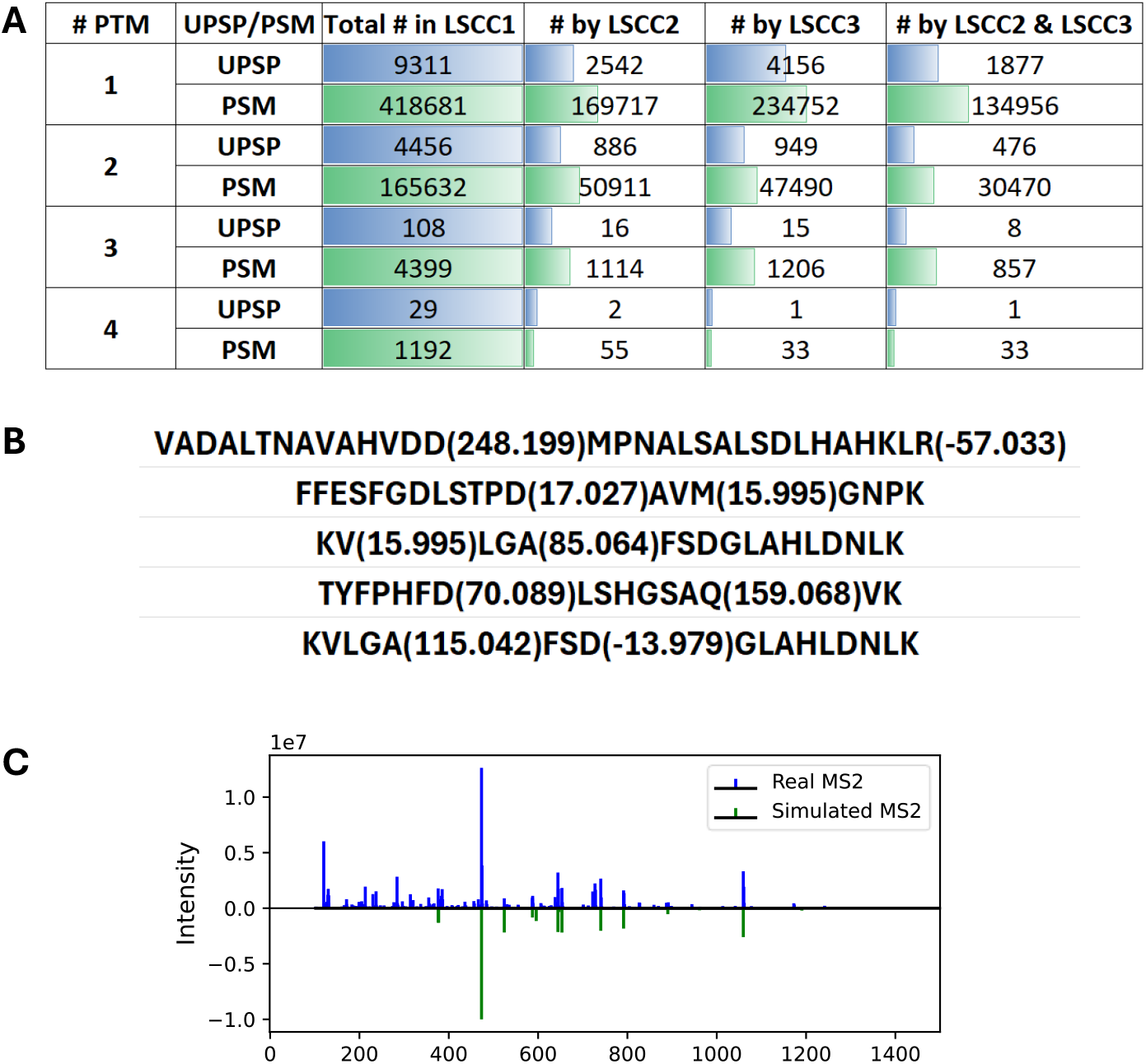
(A) The number and proportion of UPSPs and PSMs identified from the LSCC1 data that are also identified in LSCC2 and LSCC3. (B) Five examples of conserved UPSPs with two PTMs identified from LSCC1, LSCC2, and LSCC3. (C) An example of a comparison of real MS2 and simulated MS2. The real MS2 indeed contains the backbone ions with relatively high intensities.

Moreover, among the 860 upregulated UPSPs identified in LSCC1, 439 involve peptides with two PTMs, and 74 of them are also identified in both LSCC2 and LSCC3. We exemplified five of these conserved UPSP in **Fig. 8B**. These UPSPs correspond to 33 GO terms in total. 14 of these GO terms are frequently matched (more than 50 of these UPSPs per GO term), including protein binding, blood microparticle, cellular oxidant detoxification, oxygen binding, etc. In **Fig. 8C**, we simulated an MS2 that matched the second UPSP in **Fig. 8B** (F(229.163)FESFGDLSTPD(17.027)-AVM(15.995)GNPK), and compared it to one of the corresponding real MS2 (scan number 14383, fraction 13, pool 18). From this figure, we can see that the real MS2 indeed contains the backbone ions with relatively high intensities.

### 2.3 Application in COAD data

#### 2.3.1 Results of COAD1 data

We applied PIPI-C to a proteome-scale COAD data set ^95^ generated by CPTAC, denoted as COAD1. This dataset comprises 9,547,320 MS2 spectra, which were prospectively collected from 22 TMT-10 plex experiments involving 110 treatment-naive primary COAD tumors and 110 paired normal adjacent tissues. The data search took about 5 days on a 32-core server.

The peptide identification results of the COAD1 data are shown in **Fig. 9A**. At a peptide-level FDR of 0.01, we identified 5,281,641 PSMs with TMT-10 labels, among which about 44% (2,343,883) carried PTM(s) other than the TMT-10 labels, and unique modified peptides followed Zipf’s law. The distribution of all identified PTMs other than the TMT-10 labels is shown in **Fig. 9B**.

**Figure 9:**
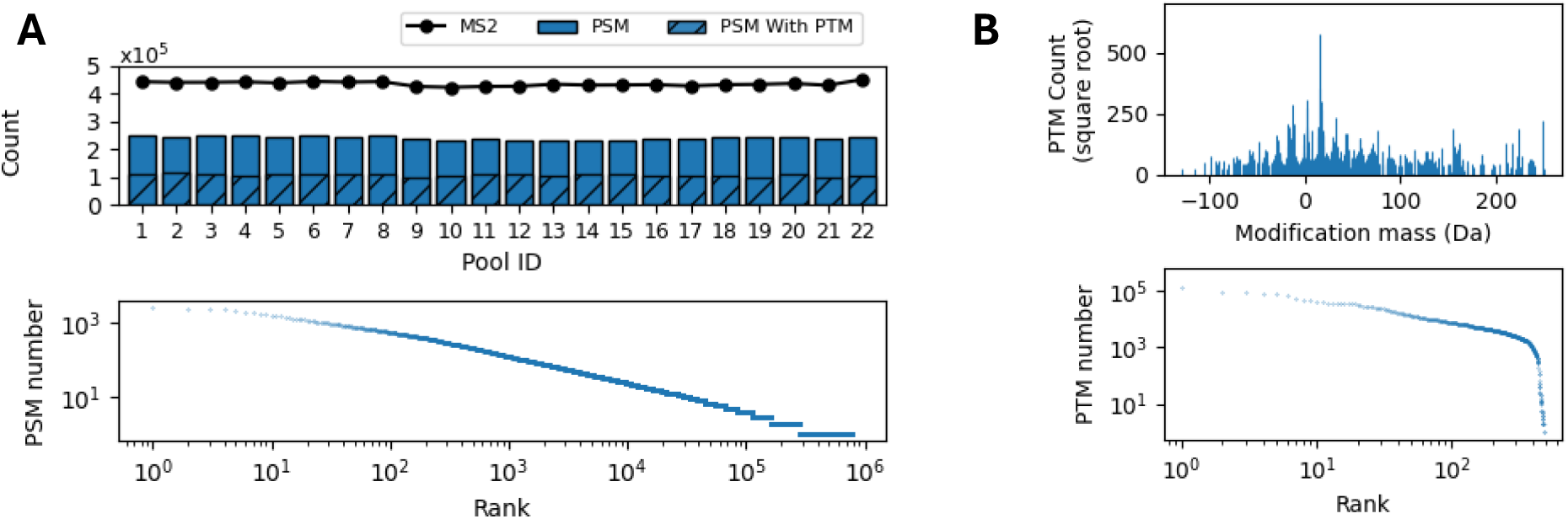
Search results of the COAD1 data using PIPI-C at a peptide-level FDR *<* 0.01. (A) Identification results: upper, the number of PSMs and those with PTMs, and the number of MS2 spectra in each experiment; lower, PSMs with differently modified peptides ranked by the PSM number. (B) PTM mass distribution with some abundant ones annotated.

The search results were then quantified, and motif analysis and GO analysis were performed. The quantification results (q-value *<* 0.05) show that 265 UPSPs have at least two PTMs other than the TMT-10 labels, corresponding to 202,264 PSMs in total (**Table 2**). The PSMs with at least one PTM were subject to quantification analysis. Among all UPSPs, 300 were significantly upregulated (q-value *<* 0.05, fold change *>* 2), and 2 were significantly downregulated (q-value *<* 0.05, fold change *<* 1/2) in the colorectal cancer sample as compared to normal samples (**Fig. 10A**). From the 300 upregulated UPSPs, we analyzed the motifs on Deamidation@N. We showed three significantly enriched motifs, namely, Gd[N], d[N]xxA, and d[N]xE with a q-value *<* 0.05 (**Fig. 10B**). The result of the GO analysis of the upregulated proteins is shown in **Fig. 10C**. We discovered significant upregulations (q-value *<* 0.01) of 31 biological processes, 74 cellular components, and 21 molecular functions, such as collagen type I trimer, IgG immunoglobulin complex, postsynaptic cytoskeleton organization, NuA4 histone acetyltransferase complex, and platelet aggregation. Some of these upregulations have been shown to be related to colon cancers. For example, an accessory subunit of human NuA4 (histone acetyltransferase), BRD8 is upregulated in human metastatic colorectal cancer and is involved in cellular survival and sensitivity to spindle poisons and proteasome inhibitors in aggressive colorectal cancers ^96,97^. Another example is that Chen et al. ^98^ found that platelet aggregation function, particularly the max aggregation ratio induced by arachidonic acid, is significantly related to the progression and clinical characteristics of colorectal cancer. Moreover, Nowak et al. ^99^ have shown that there is a distinct correlation between the metastatic capacity of human colon adenocarcinoma cells and *β*-actin (ACTB) expression, subcellular distribution, and actin cytoskeleton organization.

**Table 2:**
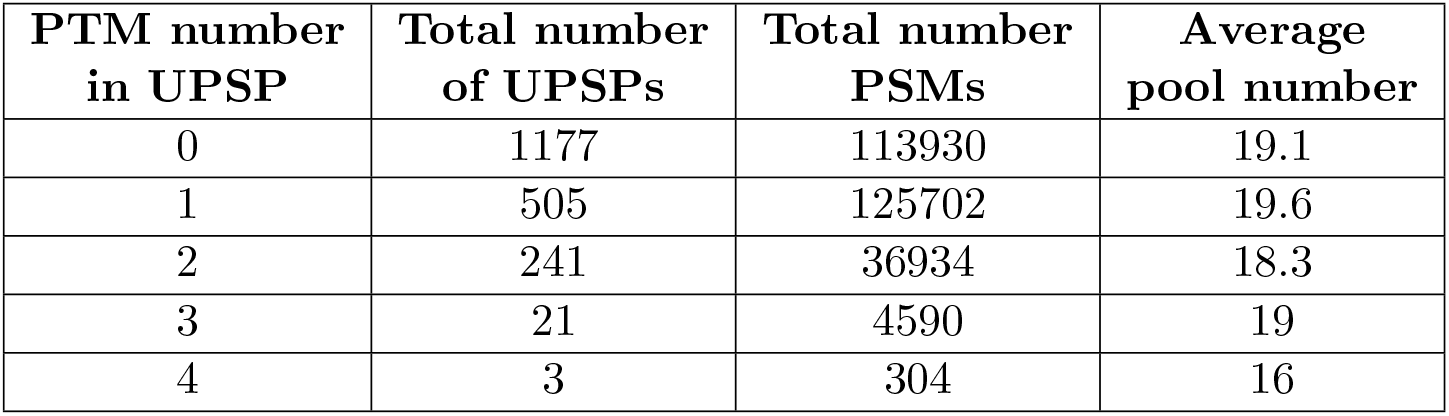
UPSPs with different numbers of PTMs identified from COAD1 data. 265 UPSPs have at least two PTMs.

| PTM number<br>in UPSP | Total number<br>of UPSPs | Total number<br>PSMs | Average<br>pool number |
| --- | --- | --- | --- |
| 0 | 1177 | 113930 | 19.1 |
| 1 | 505 | 125702 | 19.6 |
| 2 | 241 | 36934 | 18.3 |
| 3 | 21 | 4590 | 19 |
| 4 | 3 | 304 | 16 |

**Figure 10:**
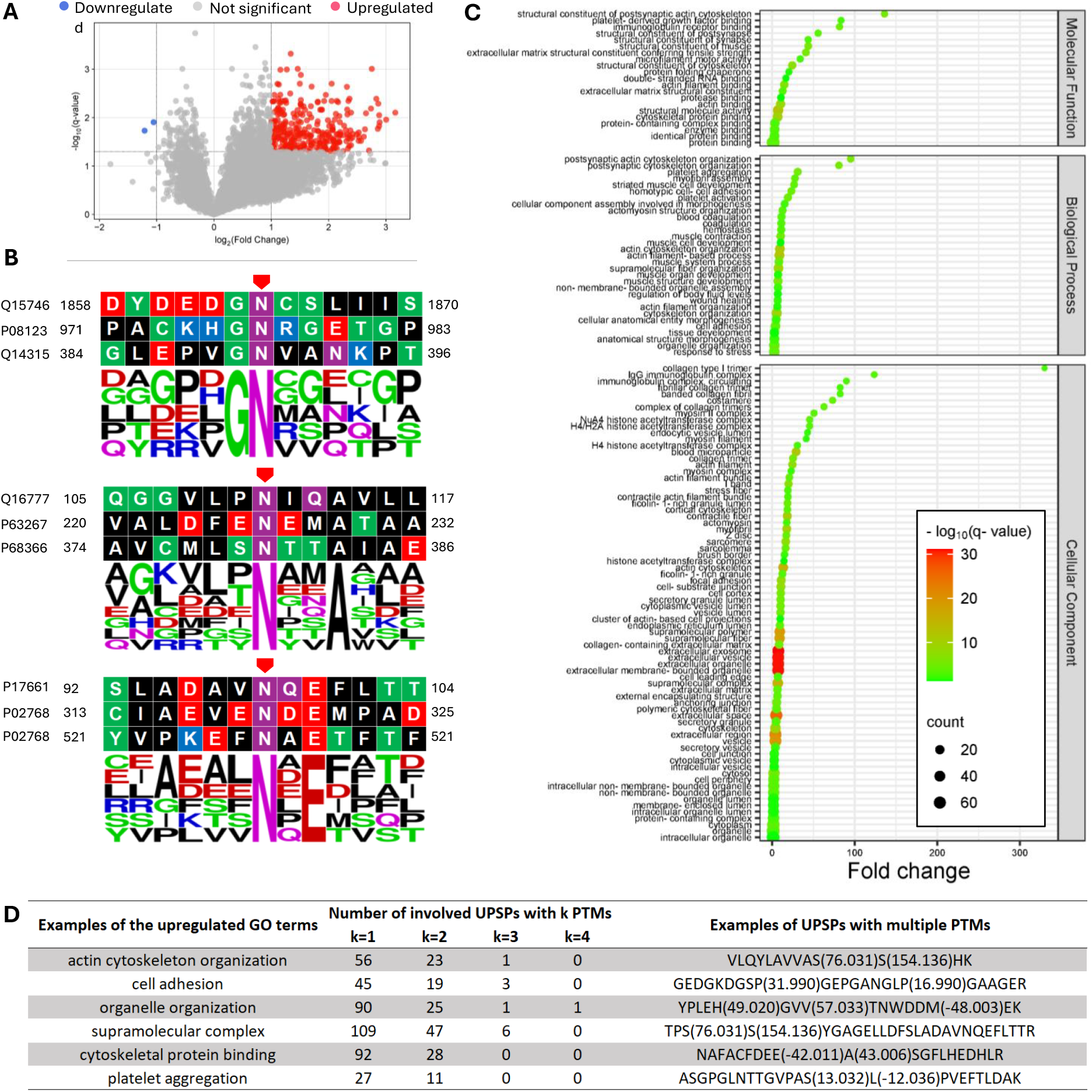
Quantification results and analysis of the LSCC data set, q-value *<* 0.05. (A) 300 upregulations (fold change *>* 2) and 2 downregulations (fold change *<* 1/2). (B) Sequence logos of upregulated Deamidation@N, q-value *<* 0.05. (C) GO analysis of the proteins with significantly regulated UPSPs. BP: biological process, BH-FDR *<* 0.01. (D) Examples of the correspondence between the upregulated GO terms and UPSPs with different numbers of PTMs.

An example of the correspondence between UPSPs and the upregulated GO terms is shown in **Fig. 10D**. About 26% (79) of the involved UPSPs contain at least two PTMs, indicating extensive potential intra-peptide PTM crosstalks in COAD1. These PTMs include hydroxymethyl, Deuterium Methylation of Lysine, tris adduct causes 104 Da addition at asparagine-succinimide intermediate, etc. Among these PTMs, 74 are reported to be related to colorectal cancer in the literature, such as acetylation ^100^, deamidation ^101^, and pursulfide ^102^. The details of all the reference papers are deposited on Zenodo at https://doi.org/10.5281/zenodo.15744650.

#### 2.3.2 Extra COAD cohort

To support the findings in the COAD1 data, we used an extra COAD cohort (COAD2) from another research group ^103^ as an independent validation. COAD2 comprises 12,618,939 MS2 spectra collected from tumors and paired distant normal tissues from 104 COAD patients. The data is labeled with TMT-10. The search parameters for this cohort are recorded in **Supplementary Table S3**.

At a peptide-level FDR of 0.01, we identified 3,182,698 with TMT labels from COAD2, among which 1,174,681 carried PTM(s) other than the TMT labels. As shown in **Fig. 11**, the first column is the number of PTMs in the peptides identified in COAD1, the third column is the total number of these peptides, and the fourth column is the number of those that are also identified in COAD2. The color bars in the cell represent their proportion in the total number of peptides (the third column). The results from COAD1 can be supported by COAD2 in two aspects. First, the rows of “UPSP” show that among UPSPs with one PTM, about 50% are identified in COAD2. Second, since some UPSPs are identified in multiple PSMs, we show the proportion of PSMs that are also identified by COAD2, i.e., about 54% of PSMs with one PTM identified in COAD1 are identified in COAD2. The proportions of PSMs with two to four PTMs identified by COAD2 are about 42%, 53%, and 5%, respectively.

**Figure 11:**
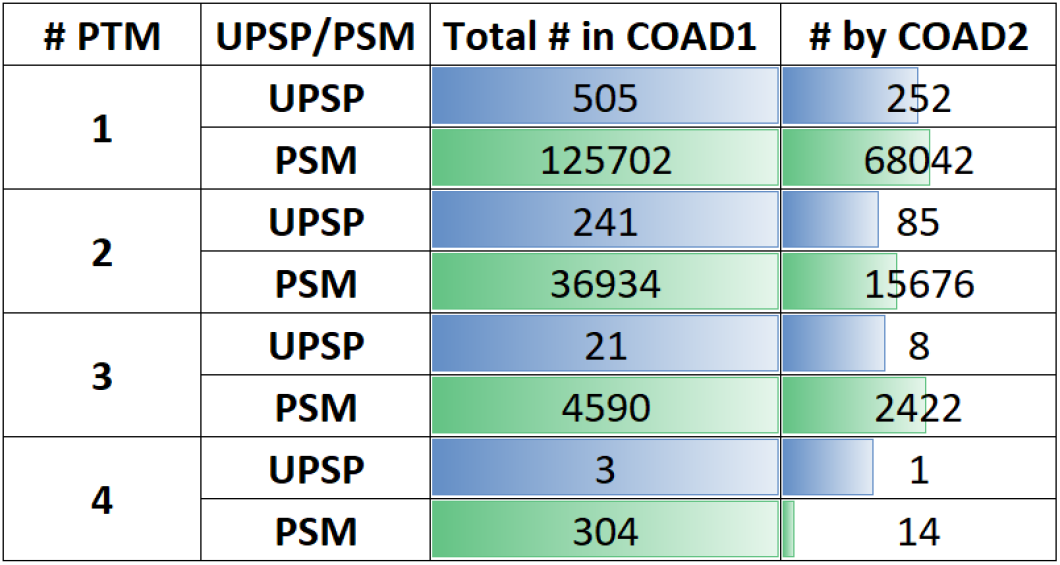
The number and proportion of UPSPs and PSMs identified from the COAD1 data that are also identified in COAD2.

### 2.4 Application in GBM data

The data sets in the LSCC and COAD application are all labeled with TMT labels. To evaluate PIPI-C’s performance on different data, we conducted this application with unlabeled data sets. We applied PIPI-C to two proteome-scale GBM cohorts (GBM1, GBM2) from different research groups. The first cohort ^104^, GBM1, comprises 7,853,872 MS2 spectra from tumor tissues of 46 GBM patients. The second cohort ^105^, GBM2, comprises 5,513,161 MS2 spectra collected from tumor tissues of 20 GBM patients. The search parameters of both cohorts are recorded in **Supplementary Table S3**. The data search took about 5 days on a 32-core server.

At a peptide-level FDR of 0.01, we identified 2,149,058 PSMs from GBM2, among which 1,167,819 carried PTM(s); these two numbers for GBM3 are 1,093,197 and 801,327. **Fig. 12A** and **Fig. 12B** shows that the ranked PSM list against the PSM numbers, the frequency number of each identified PTM, and those PTMs ranked by the identification number. We further show the interaction between the results of GBM1 and GBM2. As shown in **Fig. 12C**, the intersection of UPSPs is 23% and 19% for GBM1 and GBM2, respectively.

**Figure 12:**
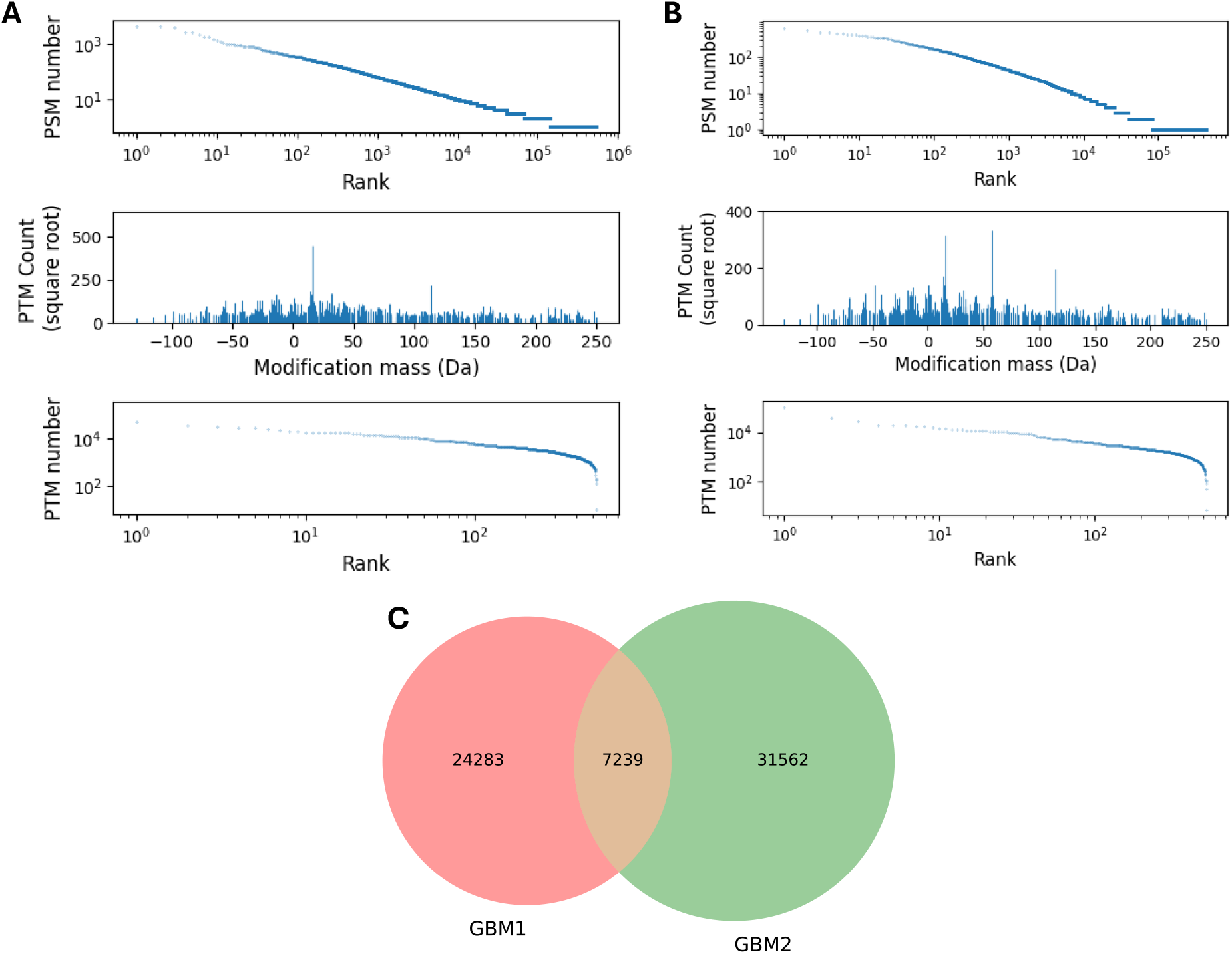
The ranked PSM list against the PSM numbers, the frequency number of each identified PTM, and those PTMs ranked by the identification number. (A) GBM1 data sets, (B) GBM2 data sets. (C) The intersection of UPSPs identified from GBM1 and GBM2.

The results show that PIPI-C is also effective from mass spectrometry data without TMT-labels.

## 3 Discussion

Due to the prevalence and significance of PTM crosstalk in peptides, identifying peptides with multiple PTMs from mass spectra data is a critical problem. This problem is essentially about finding the optimal PTM combination. Our novel search engine PIPI-C tackles this challenge by formulating the problem into an MILP model from a combinatorial optimization perspective. Notably, the backbone sequence and the PTM pattern are determined simultaneously within a single MILP model, eliminating the need for repetitive search space screening.

Through comprehensive benchmarking, PIPI-C demonstrates superior precision and sensitivity in backbone identification and PTM characterization for peptides with multiple PTMs over state-of-the-art methods. More importantly, PIPI-C can detect more PTM combinations than these existing methods, and the results were cross-validated by closed searches. In summary, the MILP model provides a theoretical formulation for the PTM characterization problem, and it can be viewed as a mathematical framework for identifying PTM crosstalk. It also allows other researchers to adopt different constraints or biological insights to modify the model formulation. PIPI-C is therefore a powerful tool for identifying peptides with multiple post-translational modifications (PTMs) without user specification, enabling large-scale research on PTM crosstalk.

In the search results of lung cancer data, we demonstrated the overlap between PIPI-C and Mascot in terms of several major PTMs in **Fig. 5**. Besides, we also performed the same analysis on the top 8 PTMs that were most frequently identified by PIPI-C, as shown in **Fig. 13**.

**Figure 13:**
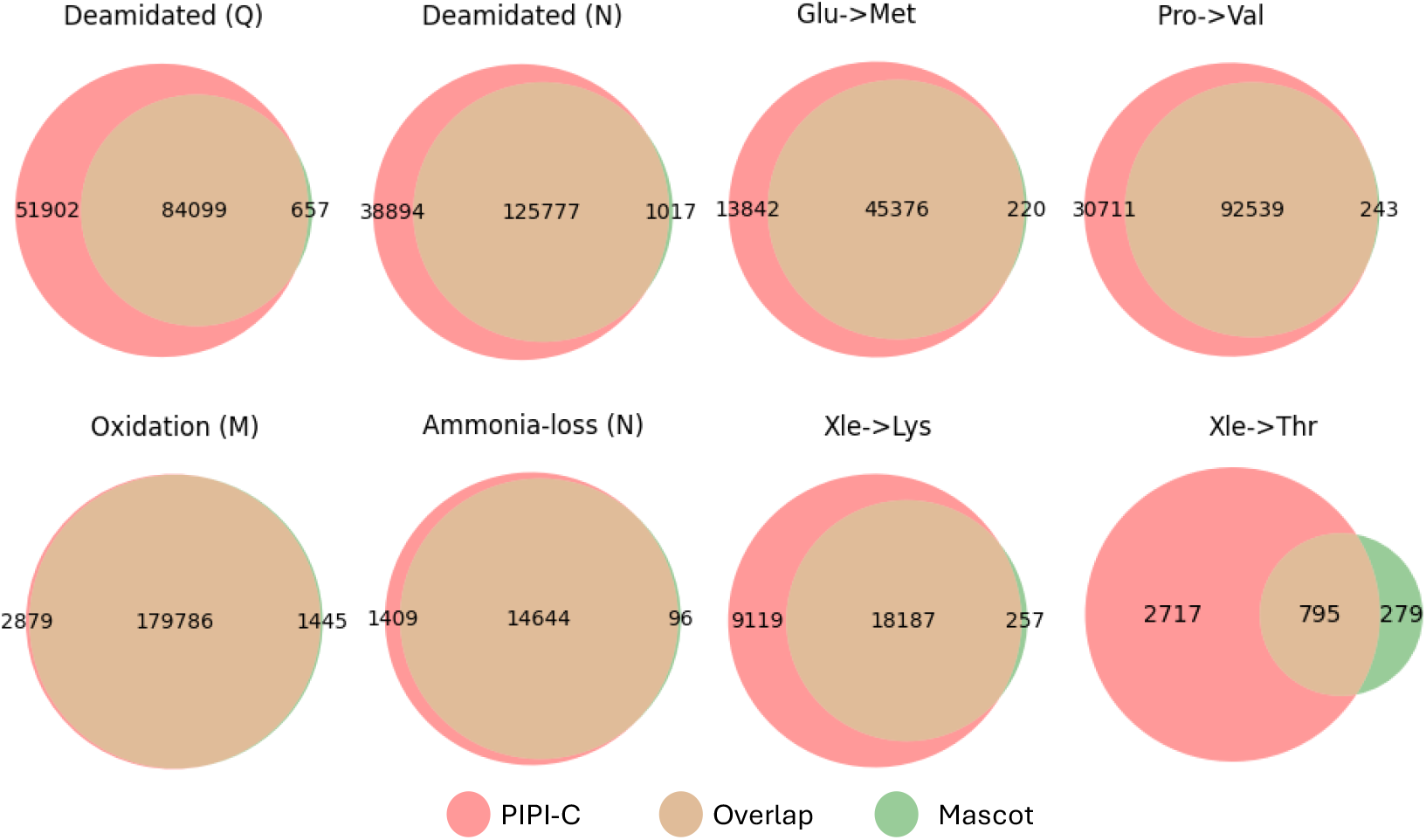
The overlap of results of PIPI-C and Mascot on the top 10 PTMs that are identified by PIPI-C.

In our analysis of the results of the lung cancer data, about 50% of the 860 upregulated UPSPs involve peptides carrying at least two PTMs. These UPSPs indicate potential PTM crosstalks in peptides. However, limited by resources, we did not conduct wet lab experiments to validate them. Instead, we conducted a thorough literature search using web-crawling techniques to find supporting evidence for the association between these PTMs and lung cancer. One supporting example is that di-methylation is related to non-small-cell lung cancer^106^. We identified potential crosstalks between di-methylation and trifluoroleucine replacement of leucine and between dimethylation and Ala to Asp substitution. Another example is that amidation is related to lung cancer ^107^, and we identified potential PTM crosstalks between amidation and Pro to Val substitution. Moreover, it is noteworthy that extracellular exosomes have a high significance level among upregulated cellular components, which are shown to be related to lung cancer in the literature. For example, Yang et al. ^108^ show that exosomes from GPC5-overexpressing lung cancer cells exert an inhibitory effect on human lymphatic endothelial cells. This outstanding significance level could potentially lead to a hypothesis for further research and biological discoveries.

While the target-decoy strategy^73^ was effectively employed to control the false discovery rate (FDR) in peptide backbone sequence identification, it was not applied to the characterization of PTMs in this paper. This decision stems from the understanding that the target-decoy approach, although well-established for peptide identification, is not designed to assess the confidence of PTM site localization or modification type assignment. To date, there is no universally accepted strategy for rigorous quality control of PTM characterization, and the development of such methods remains an open challenge in the field of proteomics. Consequently, the PTM-related results presented here should be interpreted with caution, acknowledging the potential for increased uncertainty in modification number, mass, or site assignment. Future work will benefit from integrating or developing dedicated statistical frameworks tailored to PTM validation to enhance the robustness of modification-specific findings.

Combinatorial optimization problems are inherently challenging due to the size of their exponential solution spaces, often rendering brute-force approaches computationally infeasible. To address this challenge, approximate and exact algorithms are widely adopted. Approximate methods, such as greedy algorithms, offer polynomial-time solutions (typically with time complexity ranging from linear to quadratic, depending on the implementation) but do not guarantee global optimality. These are particularly useful when scalability and responsiveness are prioritized. As exact algorithms for NP-hard combinatorial problems, MILP models are guaranteed to find the optimal solution if given sufficient time. However, their worst-case time complexity is exponential: in practice, some problem instances are complex and require extremely long solving times. To avoid unacceptable running times, we set a time limit for each case. If a case is not solved to optimality within the limit, the solver terminates, and we flag the corresponding PSM as “outOfTime”. Even if the solver stops before reaching optimality, it typically finds near-optimal solutions. This trade-off between solution quality and solving time is essential for practical applications. Based on our empirical analysis in this paper, approximately 25% of cases result in out-of-time outcomes when the time limit for each model is set to 5 seconds. One might suspect that more such cases should occur in MS2 with a larger number of PTMs. However, the proportion of out-of-time cases in MS2 with different numbers of PTMs is quite close (15% to 27%). We noticed that out-of-time cases are possible even for MS2 with only one PTM. The reason is that, given incorrect tags extracted from an MS2, we still need to build and solve MILP models, which might exceed the time limit. Thus, one possible direction for future improvement could be how to avoid spending too much time on cases with incorrect tags.

## 4 Methods

### 4.1 Overview of PIPI-C method

PIPI-C takes a protein database and raw files of MS2 spectra as input and outputs a PSM list. A raw file undergoes pre-processing using MSConverter^109^, and short amino acid sequences (called tags) are extracted from the MS2 spectra with scores equal to the total intensity of the involved peaks (**Supplementary Note 2**). These tags subsequently retrieve candidates from an FM-indexed^110^ protein database with a fuzzy and bidirectional matching approach. The mass difference between a peptide sequence and the precursor of an MS2 spectrum, denoted as Δ*m*, is considered the total mass of potential PTMs, which are then used to construct MILP models that can be solved using Gurobi. Subsequently, peptide candidates with characterized PTM patterns are collected and reranked by a protein feedback module ^64,111^. Finally, PIPI-C outputs a PSM list with FDR controlled by the target-decoy strategy.

### 4.2 Fuzzy and bidirectional match with FM-index

While many methods employ fixed-length tags to retrieve candidates from a database, PIPI-C combines tags of various lengths to obtain a better quality candidate list, thus resulting in better performance, an approach described and validated in our previous work ^64,112^. This is enabled by the FM-index, which allows efficient querying of sequences of any length in the template text. Nevertheless, it is worth noting that even a single erroneous amino acid in a tag can cause the retrieval to fail. Thus, we use a fuzzy and bidirectional matching strategy with the FM-index to accommodate erroneous amino acids at either the N-term or C-term of a tag.

The matching process is demonstrated in **Algorithm 1**. All target and decoy protein sequences are concatenated using dot signs into a long sequence, which is used to generate the original FM-indexed database (o-FMDB). The matching between a tag and the long sequence is performed in the backward direction of the tag sequence. For instance, the occurrence of a tag TAGABC is found by checking all suffixes of the tag in the following: C, CB, CBA, …, CBAGAT. If CBAGAT is finally matched, all the relative locations of TAGABC in the long sequence are output; otherwise, the retrieval fails. This situation is common, particularly with long tags that are prone to containing erroneous amino acids extracted from noise peaks. In this example, if the tag is XTAGABC, where X represents an incorrect amino acid, the matching process will fail when checking CBAGATX. As a result, the previously matched TAGABC will be discarded. To address this issue, we use a fuzzy matching strategy capable of accommodating error amino acids on the N-term of the tag. During the matching process, we record the matched amino acids and output the longest substring that matched successfully. The output for XTAGABC would be TAGABC.

However, fuzzy matching alone is insufficient where erroneous amino acids appear at the C-term of a tag, such as TAGABCX, because the initial matching is incorrect. In such cases, we employ a bidirectional match approach along with an additional reversed FM-indexed database (r-FMDB) generated from the reversed long sequence. In this approach, we attempt to match the reversed tag XCBAGAT against the r-FMDB. The backward matching process of the reversed tag is equivalent to a forward matching of the original tag. By combining this fuzzy and bidirectional matching strategy, we consistently obtain the output TAGABC, regardless of whether the erroneous amino acid X appears on the N-term or C-term of the tag, which enhances the candidate retrieval sensitivity. One might consider manually splitting the tag sequence into substrings and matching them individually to avoid generating the r-FMDB. However, such an approach would be more time-consuming. For example, if we split TAGABCX into TAGABCX, TAGABC, TAGAB, TAGA, AGABCX, GABCX, and ABCX, we would need to perform seven separate matches to the o-FMDB to find the correct one. In contrast, with the fuzzy and bidirectional matching approach, we only perform two matches: one for TAGABCX against the o-FMDB and another for XCBAGAT against the o-FMDB. Furthermore, we can generate and store the two FMDBs for a protein database and reuse them whenever needed. Compared to the overall running time, the time used for the r-FMDB generation is negligible.

#### Algorithm 1

Fuzzy and bidirectional candidate retrieval

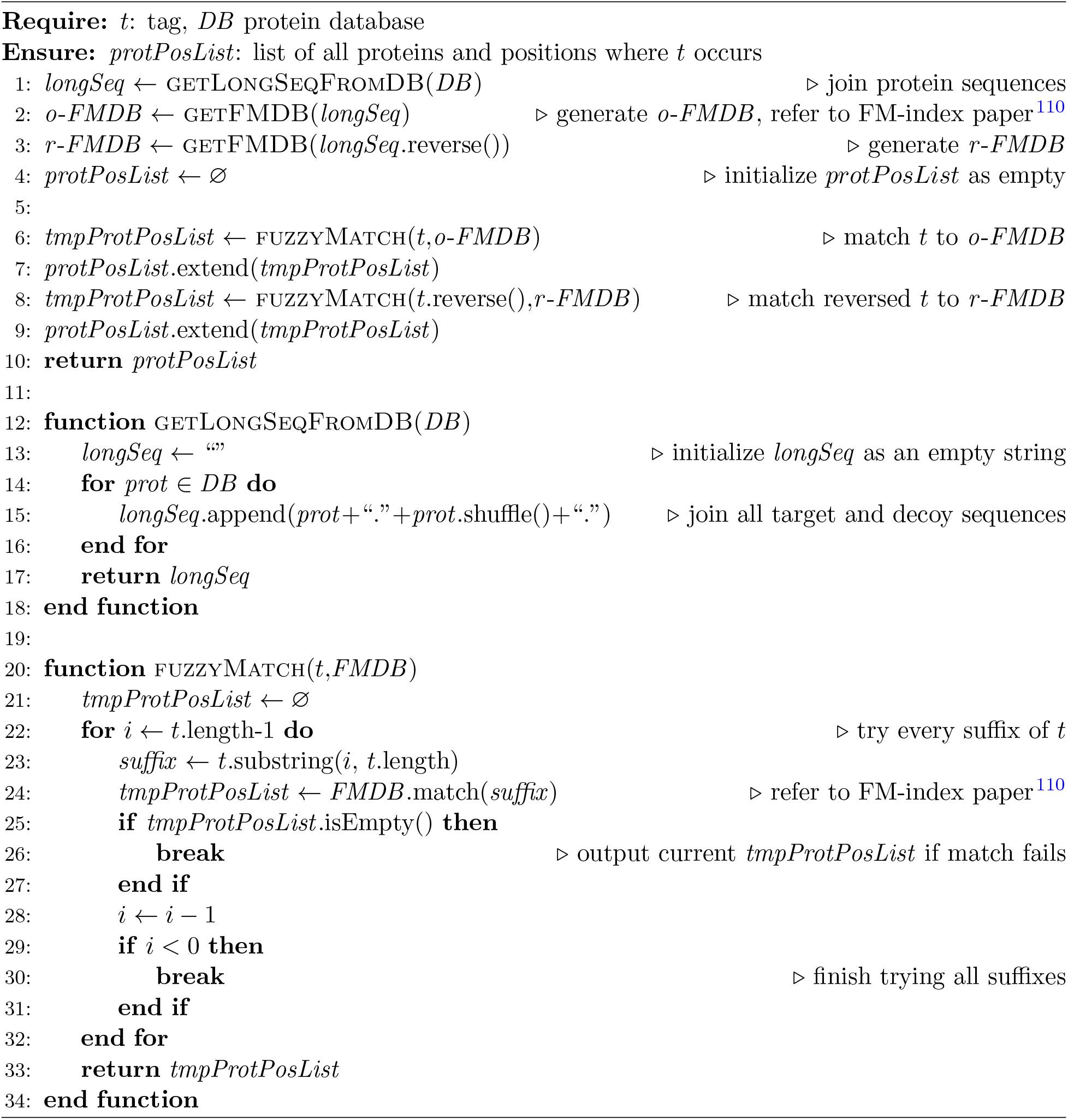

### 4.3 Protein candidates scoring

The relationship between the protein sequences and the corresponding tags located on them is a many-to-many matching. Occasionally, one protein sequence may contain multiple tags in close proximity, indicating a higher chance that the underlying peptide is digested in this region (**Fig. 14B**). Thus, we consider the proteins and either a single tag or multiple tags in close prox-imity as candidates for subsequent peptide identification and PTM characterization. Overlapping tags in close proximity are merged into one if their peaks align; otherwise, tags with lower scores are discarded as noise. Finally, the candidates are ranked by the sum of scores of the located tag(s). This aggregates the evidence from multiple tags that support the same peptide sequence, thereby avoiding redundant examination of individual tags.

**Figure 14:**
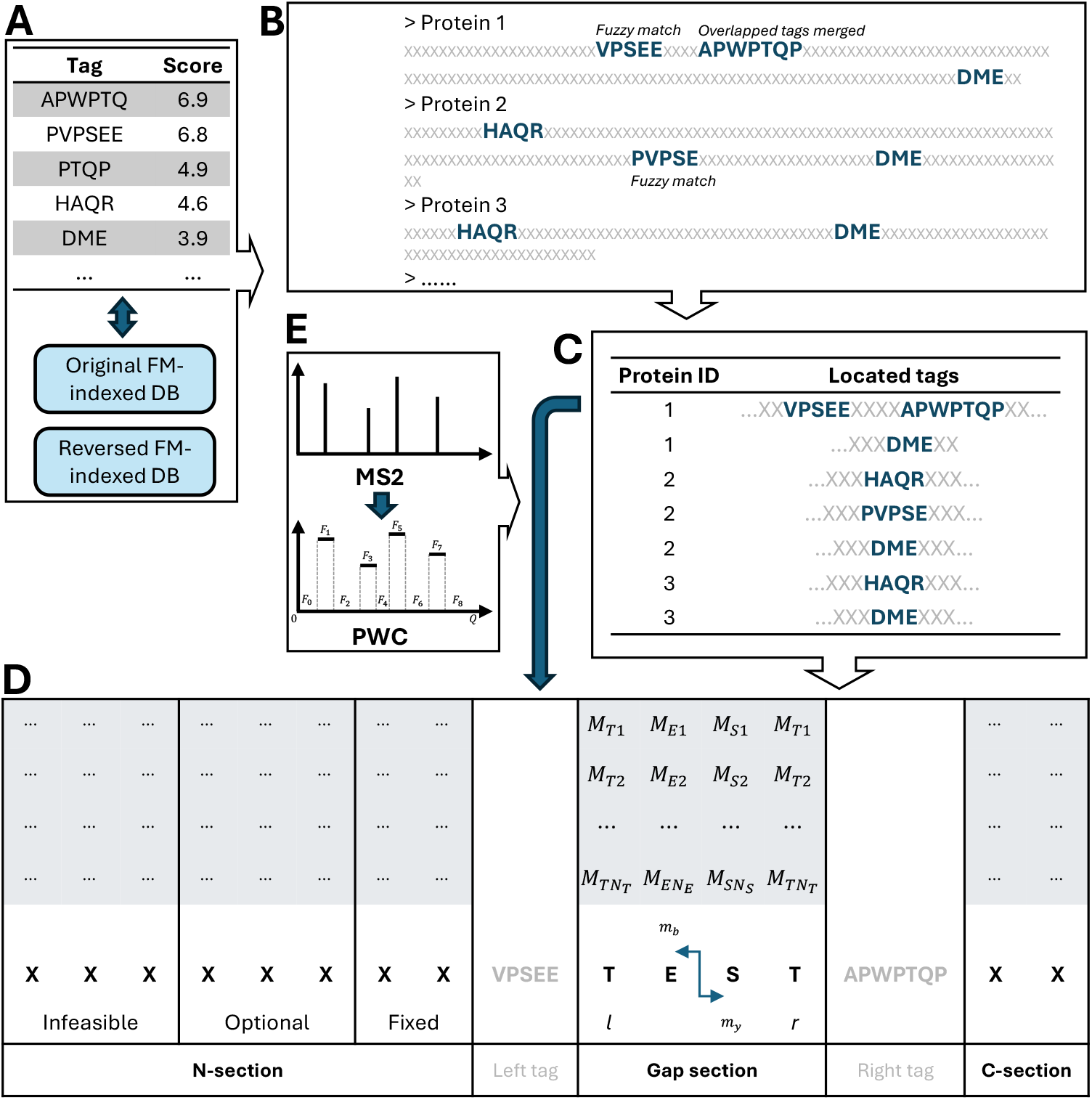
A schematic workflow of protein candidates generation. (A) Extracted tags are ranked by scores and used to retrieve protein candidates from the FM-indexed databases. (B) Tags are located on the retrieved protein sequence using a fuzzy and bidirectional matching strategy. (C) Protein sequences with located tags are considered as candidates and ranked by the total score of the located tags. (D) In either the N-section or C-section, amino acids are grouped into fixed, optional, and infeasible ones for MILP model construction. (E) The peaks of an MS2 spectrum are represented by a piece-wise constant function.

### 4.4 MILP model

Given a candidate pair, the protein sequence can be segmented into an N-section, a C-section, or gap sections by the tag(s). These sections may or may not be present in every candidate. Without loss of generality, we illustrate the MILP model using a protein sequence along with two tags that segment the protein into an N-section, a C-section, and a gap section, as shown in **Fig. 14D**. The mass shift Δ*m* in each section is known based on the peak locations of the tags, indicating that the PTM patterns in these sections are mutually independent. The assumption is that the maximal total intensity of experimental peaks, which match the theoretical peaks, is achieved when the theoretical peaks are generated from the peptide sequence with the optimal PTM pattern. The notations of the symbols are presented in Table 3.

**Table 3:**
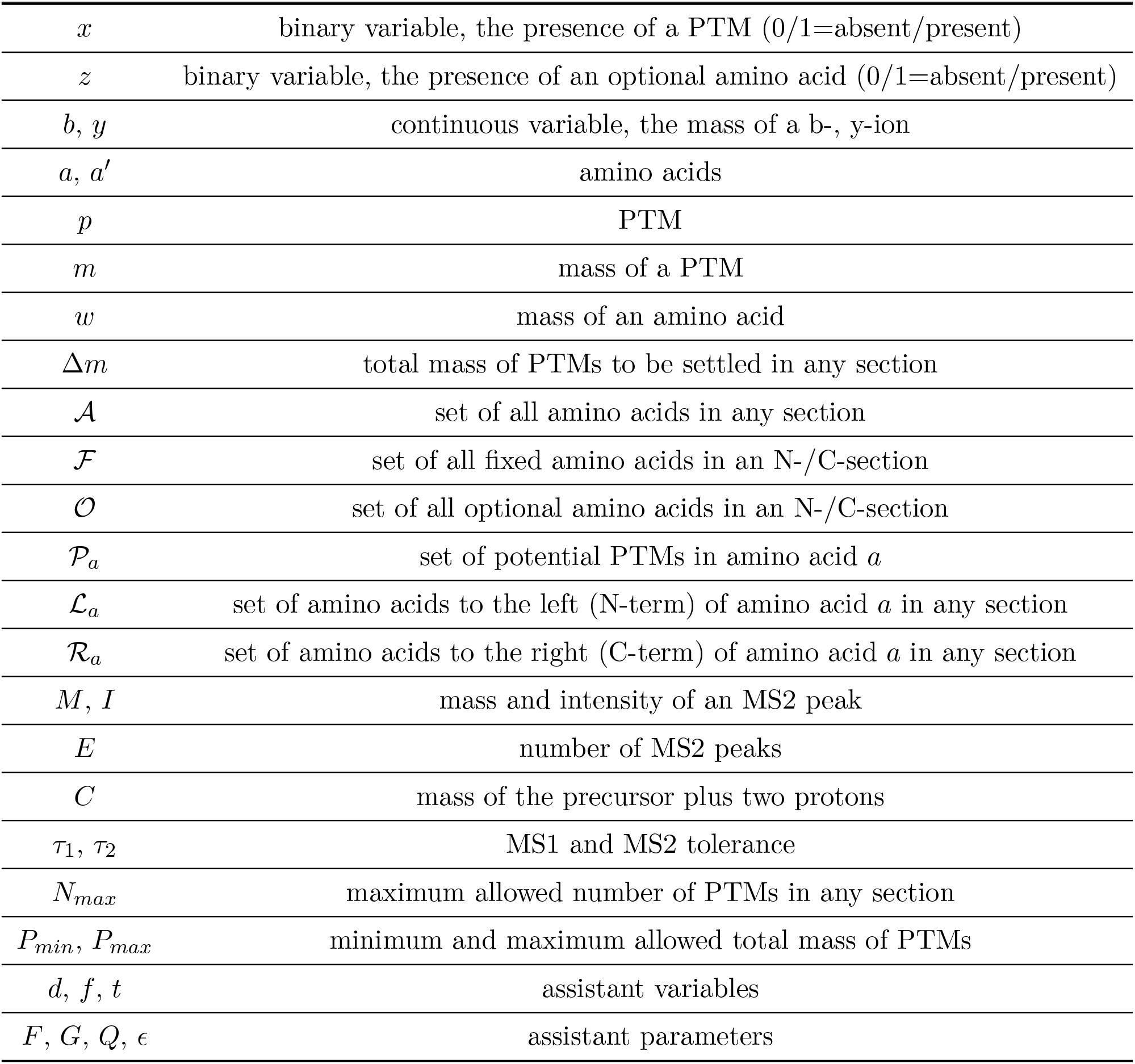
Symbols used in the MILP model.

|  |  |
| --- | --- |
| $x$ | binary variable, the presence of a PTM (0/1=absent/present) |
| $z$ | binary variable, the presence of an optional amino acid (0/1=absent/present) |
| $b, y$ | continuous variable, the mass of a b-, y-ion |
| $a, a'$ | amino acids |
| $p$ | PTM |
| $m$ | mass of a PTM |
| $w$ | mass of an amino acid |
| $\Delta m$ | total mass of PTMs to be settled in any section |
| $\mathcal{A}$ | set of all amino acids in any section |
| $\mathcal{F}$ | set of all fixed amino acids in an N-/C-section |
| $\mathcal{O}$ | set of all optional amino acids in an N-/C-section |
| $\mathcal{P}_a$ | set of potential PTMs in amino acid $a$ |
| $\mathcal{L}_a$ | set of amino acids to the left (N-term) of amino acid $a$ in any section |
| $\mathcal{R}_a$ | set of amino acids to the right (C-term) of amino acid $a$ in any section |
| $M, I$ | mass and intensity of an MS2 peak |
| $E$ | number of MS2 peaks |
| $C$ | mass of the precursor plus two protons |
| $\tau_1, \tau_2$ | MS1 and MS2 tolerance |
| $N_{max}$ | maximum allowed number of PTMs in any section |
| $P_{min}, P_{max}$ | minimum and maximum allowed total mass of PTMs |
| $d, f, t$ | assistant variables |
| $F, G, Q, \epsilon$ | assistant parameters |

#### 4.4.1 MILP for the gap section

We start with the simpler case within the gap between a left and right tag. Δ*m* is computed as the mass difference between the start peak of the right tag and the end peak of the left tag, i.e., Δ*m* = *M*_*r,start*_ −*M*_*l,end*_. The set of amino acids within the gap is denoted as 𝒜. The set of potential PTMs on amino acid *a* is denoted as *𝒫_a_*. For each *p* ∈ *𝒫_a_*, where *a* ∈ *𝒜*, a binary variable *x* is assigned to indicate the presence of *p*. The masses of b- and y-ions, *b* and *y*, are expressed as linear combinations of *x* and other constants. The objective is to maximize the total intensity of the matched experimental peaks by all b- and y-ions. The model is formulated as follows:

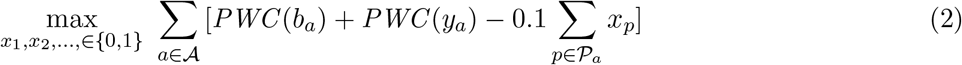

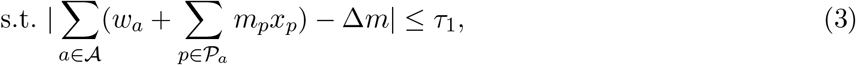

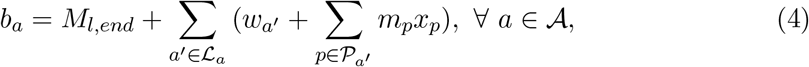

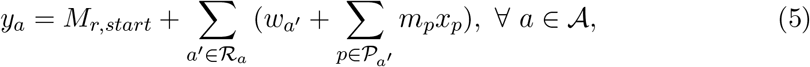

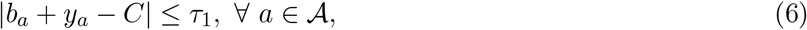

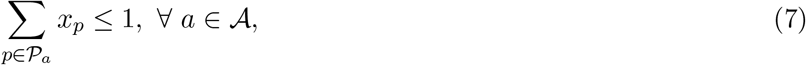

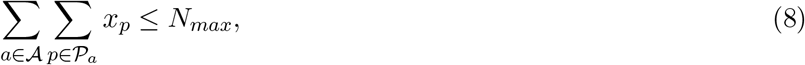

where | *·* | is the absolute value of a scalar or the cardinality of a set, and ℒ_*a*_ and ℛ_*a*_ are the sets of amino acids to the left or right of amino acid *a*. Constraint (3) enforces that the difference between the total mass of PTMs and Δ*m* is within *τ*_1_; constraint (4) and (5) express *b* and *y* with PTM masses, amino acid masses, and peaks of the left and right tags; constraint (6) indicates the relationship between each pair of *b* and *y* at each *a*; constraint (7) enforces that at most one PTM can occur on each amino acid; constraint (8) is an upper limit of the total number of PTMs. The objective function is the summation of the intensities of peaks matched by b- and y-ions, plus a penalty term on the total number of PTMs. *PWC* (*·*) is a piece-wise constant (PWC) function that represents the MS2 peak intensities and zeros between two MS2 peaks. As shown in **Fig. 14E**, these values are denoted as (*F*_0_, *F*_1_, …, *F*_2*E*_), where *E* is the number of MS2 peaks. Thus we have

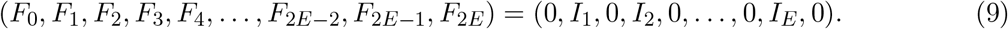

As shown in equation (10), if the input mass *t* matches the mass of any peak *M*_*e*_, the function returns the corresponding intensity *I*_*e*_; otherwise, it returns 0.

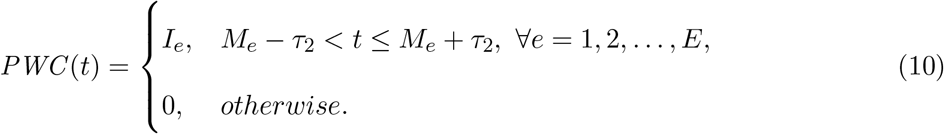

*PWC* (*·*) is not inherently linear but can be linearized using auxiliary variables. We define binary variables (*d*_0_, *d*_1_, …, *d*_2*E*_), one for each *F*_*i*_, and a continuous variable *f* for the return value of *PWC* (*t*). Then, we apply the constraints as follows:

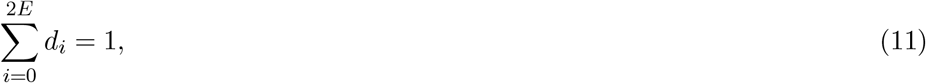

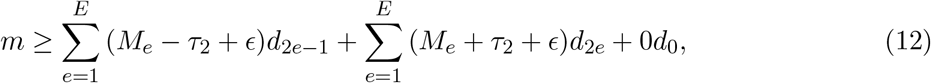

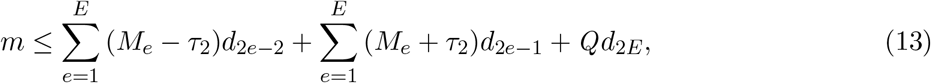

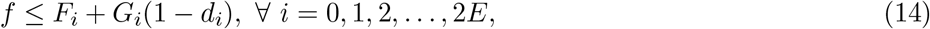

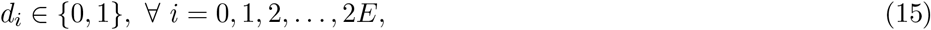

where *Q* is a constant that upper bounds all *M*_*e*_, *G*_*i*_ is a big constant such that *F*_*i*_ +*G*_*i*_ ≥ *F*_*j*_, *∀j* ≠ *i*, and *ϵ* is a small tolerance constant. Constraint (11) enforces that only one *F*_*i*_ can be selected; constraint (12) and (13) enforce that *M*_*i*−1_ *≤ m ≤ M*_*i*_ *⇒ d*_*i*_ = 1; constraint (14) imposes an upper bound on the return value *f*, restricting it to be *F*_*i*_ only when *d*_*i*_ = 1 because other inequalities where *d*_*i*_ = 0 become redundant because of *G*_*i*_. Since the objective function is maximum-wise, *f* will automatically assume the value of *F*_*i*_. As a result, the objective function can be equivalently replaced with

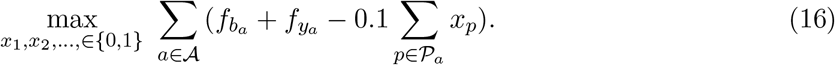

#### 4.4.2 MILP for the N- and C-section

The models for the N- and C-sections are similar. Thus, without loss of generality, we only present the one for the N-section here. Taking the first candidate in **Fig. 14C** as an example, VPSEE is the right tag for the N-section, and no left tag is available. The N-section ends at the right tag, but the starting amino acid site is unknown without a left tag, i.e., the peptide sequence needs to be determined. To achieve this, we could extend the N-section towards the protein N-term, building and solving one MILP model at every potential position, which is quite time-consuming and involves repetitive work. Instead, we incorporate the extension step into a single model, which simultaneously enables the determination of the peptide sequence and the characterization of the PTM pattern.

In the N-section, Δ*m* equals the start peak of the right tag 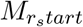. Given *P*_*min*_ and *P*_*max*_, the amino acids on the N-term side of the right tag can be classified into fixed, optional, and infeasible categories. From the right tag towards the protein N-term, the first several amino acids are in the fixed amino acids set ℱ; without these amino acids, the total mass of present PTMs will be beyond *P*_*max*_. The following several amino acids are in the optional amino acids set 𝒪, which may or may not be present in the sequence. The remaining amino acids are infeasible, with which the total mass of present PTMs will be smaller than *P*_*min*_. In summary, the amino acids in ℱ and 𝒪 are selected and added to the sets from the ones in the N-section in the direction from the right tag to the N-term, satisfying the following conditions:

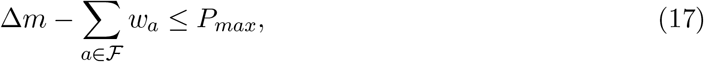

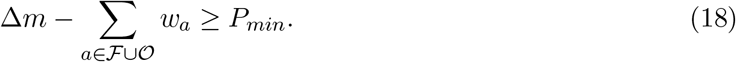

*b*_*a*_ and *y*_*a*_ are calculated similarly to those in the gap section, except that *M*_*l,end*_ = 0 for the N-section.

Then, the model is written as follows:

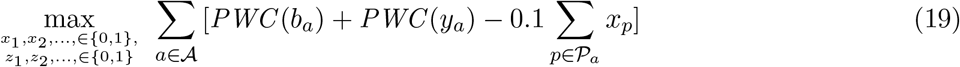

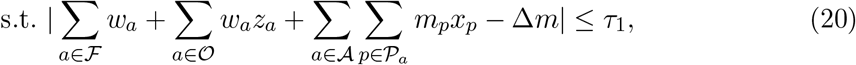

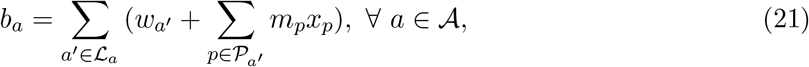

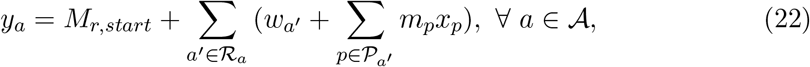

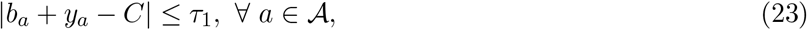

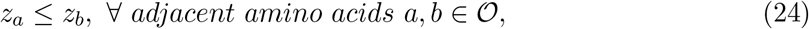

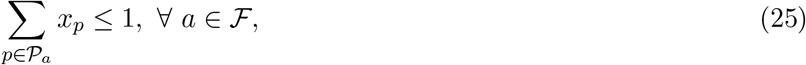

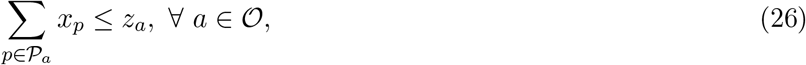

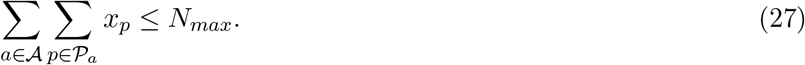

he objective function (19) can be linearized using the same approach as in the gap section. The differences between the model for the gap section and the one here are as follows: for *a* ∈ 𝒪, the mass is only added when *z*_*a*_ = 1 as constrained by (20); *M*_*l,end*_ in constraint (21) is omitted for being zero; constraint (24) enforces that, for every two adjacent optional amino acids, the left one can be present only if the right one is present; and constraint (26) enforces that if an optional amino acid is absent, no PTM can occur on it.

### 4.5 The three cancer data applications

We applied PIPI-C to uncover the PTM profile underlying the LSCC, COAD, and GBM data. The TMT label was set as a variable modification on the peptide N-term and all Serines, and set as a fixed modification on all Lysines. Full parameter settings are recorded in **Supplementary Table S3**. Original parameter files are deposited on Zenodo at https://doi.org/10.5281/zenodo.15744650.

#### 4.5.1 Quantification

PSMs without a TMT label were excluded from the analysis. Peptides with more than one PSM (PSM number *>* 1) were selected as quantifiable. The intensity of the TMT reporter ion for each PSM was extracted from the original mgf data. It was assumed that each sample would contain a set of unregulated peptides with abundances comparable to the common reference (CR) sample, meaning these peptides should have a log TMT ratio centered at zero in the normalized sample. Consequently, two-component normalization of each TMT label’s intensity was applied against the CR^77^.

PSM-level log ratios were calculated as the mean log values for each patient within the corresponding PSM. Peptides containing the same PTM sites were grouped into a UPSP, and the UPSP ratio was calculated as the mean value of all contributing PSM-level ratios. Each pool of the dataset was treated as an individual experiment, and a UPSP was required to have different mean log ratios in each pool. Only UPSPs present in more than 60% of the pools were selected for further quantification. A one-sample t-test was conducted to generate the p-value for each UPSP, followed by BH-FDR (Benjamini-Hochberg FDR) correction to produce the q-value.

### 4.5.2 Motif analysis

Motifs were extracted by Motif-X ^113^ via the MEME suite ^114^, and plotted by WebLogo ^115^. The foreground data set was obtained from all the peptides with Phosphorylation@S, with fold change *>* 3. In total, 43 sequences were input for motif discovery. These peptide sequences were centered on Phosphorylation@S and extended to 13 amino acids (*±*6 on both the peptide N- and C-term). When the extension was unavailable on the term, “X” would be added to keep the same sequence length. Background peptides were extracted from all peptide sequences of length 13 in the Uniprot Human protein database, which were centered on S. The minimum number of motif occurrences was set to 8, and the significance threshold was set to q-value *<* 0.01.

#### 4.5.3 Gene ontology enrichment analysis

All proteins that contain quantified and modified peptides (q-value *<* 0.05 and fold change *>* 3) were further analyzed for GO enrichment against the human genome as the background data set. The statistical significance of GO analysis was calculated by a two-tailed Fisher exact test and followed by BH-FDR correction. GO terms with q-value *<* 0.01 were output and plotted.

#### 4.5.4 Interactome analysis

We incorporated PPI data sourced from the STRING database^116^, which aggregates interaction information from various sources such as text mining, experiments, databases, co-expression, neighborhood, gene fusion, and co-occurrence. The PPI network was visualized using Cytoscape (version 3.10.2) ^117^. In this network representation, each pair of interacting proteins and the link between them were denoted as nodes and edges.

## Data availability

All parameter files, result files, and the simulation data sets are deposited on Zenodo at https://doi.org/10.5281/zenodo.15744650. The synthetic data sets, soybean data sets, *Petunia* data sets, LSCC2, GBM1, and GBM2 data sets can be downloaded from the ProteomoXchange Consortium with the data identifiers PXD009449, PXD034796, PXD005470, PXD010429, PXD016165, and PXD019381, respectively. The LSCC1 and COAD1 data sets can be downloaded from the Proteomic Data Commons^118^ with the data identifiers PDC000234 and PDC000116, respectively. The LSCC3 and COAD2 data sets can be downloaded from the iProx database (Integrated Proteome Resources) (http://www.iprox.org/index) using the data identifiers IPX0001833000 and IPX0005732001, respectively.

## Code availability

PIPI-C was developed in the Java programming language. The source code can be accessed on GitHub at https://github.com/lsz1994024/PIPI-C and https://bioinformatics.hkust.edu.hk/Software/PIPI-C.html.

## Acknowledgements

The study was partially supported by grants from the Hong Kong Research Grants Council (RGC): 16102422, 16103621, T12-101/23-N, C7015-23G, and R4012-18; MHP/033/20 and ITS/043/23 from the Innovation and Technology Commission (ITC) of the Hong Kong S.A.R., the Hetao Shenzhen-Hong Kong Science and Technology Innovation Cooperation Zone project (HZQB-KCZYB-2020083), and internal grants 3030 009, BGF.001.2023, Z1056, and CSSET24SC01 from HKUST.

## Author contributions

S.L. designed the algorithms, developed PIPI-C software, and designed the validation experiments; S.L., P.Z., and S.D. performed the experiments; S.L., S.D., P.Z, and C.Z. performed the validation experiments and analyzed the LSCC, COAD, and GBM data; S.L. wrote the manuscript; N.L. and W.Y. supervised this study. All authors have read and approved this manuscript.

## Additional information

### Supporting information

The paper contains supplementary notes, figures, and tables discussing the issues below. Supplementary Tables 5-9 are large and placed in the Excel file named “Supplementary Tables.xlsx”, which is deposited on Zenodo at https://doi.org/10.5281/zenodo.15744650.

**Supplemental Note** 1: Generation of simulated data sets

Table S1. Parameters, PTMs, and sites used in AlphaPeptDeep for simulated data sets.

**Supplemental Note** 2: Data preprocessing and tag extraction **Supplemental Note** 3: Software and parameters used in the experiments Table S2. Versions of software programs used in this work.

Table S3. Common parameters used in the experiments.

**Supplemental Note** 4: Details of the 21 synthetic data sets and the results

Table S4. Information of the 21 synthetic data sets. Each data set contains tens of thousands of MS2 spectra with up to one PTM.

Fig. S1. Comparison results of the synthetic data sets 01 to 07 at FDR of 0.01. Fig. S2. Comparison results of the synthetic data sets 08 to 14 at FDR of 0.01. Fig. S3. Comparison results of the synthetic data sets 15 to 21 at FDR of 0.01. **Supplemental Note** 5: Results of replicated soybean data set.

Fig. S4. Replicate R02. Intersections of PSMs with fully dimethyl-labeled peptides identified from the soybean data sets.

Fig. S5. Replicate R03. Intersections of PSMs with fully dimethyl-labeled peptides identified from the soybean data sets.

Fig. S6. Replicate R05. Intersections of PSMs with fully dimethyl-labeled peptides identified from the soybean data sets.

Fig. S7. Replicate R05. Intersections of PSMs with fully dimethyl-labeled peptides identified from the soybean data sets.

Fig. S8. Replicate R06. Intersections of PSMs with fully dimethyl-labeled peptides identified from the soybean data sets.

**Supplemental Note** 6: More results of the *Petunia* data set.

Fig. S9. For Open-pFind and MODplus, unidentified PTM combinations can also be identified when the other PTM is pre-specified as variable modification.

## Correspondence

Correspondence and material requests should be addressed to Ning Li or Weichuan Yu.

