## Supplementary Information for "PIPI-C: A combinatorial optimization framework for identifying post-translational modification crosstalks in mass spectrometry data"

for

### Contents


**Contents** **S2**

**1 Supplementary Note 1: Generation of simulated data sets** **S4**

Table S1. Parameters, PTMs, and sites used in AlphaPeptDeep for simulated data sets S4

**2 Supplementary Note 2: Data preprocessing and tag extraction** **S5**

**3 Supplementary Note 3: Software and parameters used in the experiments** **S6**

Table S2. Versions of software programs used in this work . . . . . . . . . . . . . . . . S6

Table S3. Common parameters used in the experiments. A single asterisk (*) denotes the search engine incorporating the corresponding variable modification setting. Double asterisk (**) denotes the search engine with both Phe→Cys@F and Unknown:248@N-term as variable modifications. . . . . . . . . . . . . . . S7

**4 Supplementary Note 4: Details of the 21 synthetic data sets and the results** **S8**

Table S4. Information of the 21 synthetic data sets. Each data set contains tens of thousands of MS2 spectra with up to one PTM. . . . . . . . . . . . . . . . . . . S8

Figure S1. Comparison results of the synthetic data sets 01 to 07 at FDR of 0.01: bar chart, PSM numbers; pie chart, PTM numbers. . . . . . . . . . . . . . . . . . . S9

Figure S2. Comparison results of the synthetic data sets 08 to 14 at FDR of 0.01: bar chart, PSM numbers; pie chart, PTM numbers. . . . . . . . . . . . . . . . . . . S9

Figure S3. Comparison results of the synthetic data sets 15 to 21 at FDR of 0.01: bar chart, PSM numbers; pie chart, PTM numbers. . . . . . . . . . . . . . . . . . . S10

**5 Supplementary Note 5: Results of replicated soybean data set** **S11**

Figure S4. Replicate R02. Intersections of PSMs with fully dimethyl-labeled peptides identified from the soybean data sets. D1, D2: two samples in the data set. S11

Figure S5. Replicate R03. Intersections of PSMs with fully dimethyl-labeled peptides identified from the soybean data sets. D1, D2: two samples in the data set. S12

Figure S6. Replicate R04. Intersections of PSMs with fully dimethyl-labeled peptides identified from the soybean data sets. D1, D2: two samples in the data set. S12

Figure S7. Replicate R05. Intersections of PSMs with fully dimethyl-labeled peptides identified from the soybean data sets. D1, D2: two samples in the data set. S13

Figure S8. Replicate R06. Intersections of PSMs with fully dimethyl-labeled peptides identified from the soybean data sets. D1, D2: two samples in the data set. S13

**6 Supplementary Note 6: More results of the *Petunia* data set S14**

Figure S9. For Open-pFind and MODplus, unidentified PTM combinations can also be identified when the other PTM is pre-specified as variable modification . . . S14

### 1 Supplementary Note 1: Generation of simulated data sets

We used AlphaPeptDeep[1] to generate simulated MS2 spectra with selected PTMs as shown in **Table S1**. We allowed up to four PTMs in one peptide. Other parameters include a maximum missed cleavage site of 2. Since AlphaPeptDeep only predicted $b$ and $y$ ions, we manually added noise peaks to generate full MS2 spectra. The signal-to-noise ratio (SNR) is controlled by

$$\mathrm{SNR} = \sum I_s^2 / \sum I_n^2, \tag{S1}$$

where subscript $s$ and $n$ denote signal and noise, respectively. We obtained seven data sets from 881 template peptide sequences with different average SNRs ranging from 2.023 to 0.79. This SNR range was set based on the data referenced in the literature[2–4].

Table S1: Parameters, PTMs, and sites used in AlphaPeptDeep for simulated data sets

| | | | |
|---|---|---|---|
| **Enzyme** | trypsin | | Oxidation@G |
| **Instrument** | Lumos | | Carboxyethyl@H |
| **Max variable modifications** | 4 | | Carbonyl@L |
| **Max missed cleavages** | 2 | | Oxidation@M |
| **Min precursor charge** | 2 | | Deamidated@N |
| **Max precursor charge** | 4 | | Dioxidation@P |
| **Min peptide length** | 8 | **Variable modifications** | Deoxyhypusine@Q |
| **Max peptide length** | 25 | | Ethanolyl@R |
| **Fix modification** | Carbamidomethyl@C | | Malonyl@S |
| **Variable modifications** | Fluoro@A | | Methylamine@T |
| | Sulfide@D | | Carbonyl@V |
| | Decarboxylation@E | | Chlorination@W |
| | Nitro@F | | Phospho@Y |

#### 2 Supplementary Note 2: Data preprocessing and tag extraction

The raw file is first converted into mgf format using ProteomeWizard MSConvert[5], then decharged and deisotoped using MS2-Deisotoper[6]. For each MS2 spectrum, the entire $m/z$ range is divided into subranges of 100 Daltons, and the intensities of peaks are normalized by dividing them by the highest intensities in the corresponding subranges.

A weighted directed graph $G$ is then constructed for tag extraction. The $m/z$ difference between any two peaks is calculated to match potential amino acids. When an amino acid is matched, two nodes, $n_i$ weighted by $I_i$ (intensity of peak $i$), $n_j$ weighted by $I_j$ (intensity of peak $j$), and a directed edge $e_{i,j}$ weighted by $I_i + I_j$ are added to $G$. All paths from nodes with a zero in-degree to nodes with a zero out-degree are extracted as tags using depth-first search and ranked by the sum of the weights of nodes involved. All tags longer than the minimum tag length defined in the parameter file are collected.

### 3 Supplementary Note 3: Software and parameters used in the experiments

The version of all software used in this work is shown in **Table S2**. The common parameters used for the search engines in all the experiments are recorded in **Table S3**. All full parameter files are deposited on Zenodo at https://doi.org/10.5281/zenodo.14885715.

Table S2: Versions of software programs used in this work

| | |
|---|---|
| Open-pFind[7] | 3.2.0 |
| MODplus[8] | 2.01 |
| Gurobi (Academic license)[9] | 10.0.3 |
| MSConvertGUI[5] | 3.0.22238 |
| MS2-Deisotoper[6] | N/A |
| MEME Suite[10] | 5.5.5 |
| Mascot[11] | 2.7.0 |

Table S3: Common parameters used in the experiments. A single asterisk (*) denotes the search engine incorporating the corresponding variable modification setting. Double asterisk (**) denotes the search engine with both Phe→Cys@F and Unknown:248@N-term as variable modifications.

| **Exp** | **Software** | **MS1 tolerance** | **MS2 tolerance** | Max cleavage sites | Fixed mod | Variable mod | Enzyme |
|---|---|---|---|---|---|---|---|
| 1 | PIPI-C | 10ppm | 0.01 Da | 2 | Carbamidomethyl@C | Oxidation@M | Trypsin |
| | Open-pFind | | | | | | |
| | MODplus | | | N/A | | | |
| 2 | PIPI-C | | | 1 | | | |
| | Open-pFind | | | | | | |
| | MODplus | | | N/A | | | |
| 3 | PIPI-C | | | 1 | | | |
| | Open-pFind | | | | | | |
| | MODplus | | | N/A | | | |
| 4 | PIPI-C | | 0.02 Da | 2 | | Oxidation@M Acetyl@Prot-NTerm | |
| | Open-pFind | | | | | | |
| | MODplus | | | N/A | | | |
| | Open-pFind* | | | 2 | | Oxidation@M Acetyl@Prot-Nterm, one undetected PTM | |
| | MODplus* | | | N/A | | | |
| | Mascot | | | 2 | | Oxidation@M Acetyl@Prot-Nterm, one undetected PTM, GG@K | |
| | Open-pFind** | | | | | Oxidation@M Acetyl@Prot-Nterm, one undetected PTM, Phe->Cys@F, Unknown248@AnyNterm | |
| LSCC1 | PIPI-C | | 0.01 Da | | Carbamidomethyl@C TMT@K | TMT@Pep-N-term, TMT@S | |
| LSCC2 | PIPI-C | | 0.02 Da | | | | |
| LSCC3 | PIPI-C | | 0.02 Da | | | | |
| COAD1 | PIPI-C | | 0.01 Da | | | | |
| COAD2 | PIPI-C | | 0.02 Da | | | | |
| GBM1 | PIPI-C | | 0.02 Da | | Carbamidomethyl@C | Oxidation@M | |
| GBM2 | PIPI-C | | 0.02 Da | | | | |

### 4 Supplementary Note 4: Details of the 21 synthetic data sets and the results

The details of all the 21 synthetic data sets[12] including the one used in experiment 2 are collected in Table S4. All the results are shown in **Figure S1**, **Figure S2**, and **Figure S3**.

Table S4: Information of the 21 synthetic data sets. Each data set contains tens of thousands of MS2 spectra with up to one PTM.

| ID | Residue | Modification | Monoisotopic mass | Number of MS2 |
|---|---|---|---|---|
| 1 | Lysine | Formylation | 27.995 | 51861 |
| 2 | Lysine | Acetylation | 42.010 | 50511 |
| 3 | Tyrosine | Phosphorylation | 79.966 | 55131 |
| 4 | Lysine | Methylation | 14.016 | 49086 |
| 5 | Lysine | Biotinylation | 226.078 | 43572 |
| 6 | Lysine | Butyrylation | 70.042 | 48573 |
| 7 | Lysine | Crotonylation | 68.026 | 49398 |
| 8 | Lysine | Dimethylation | 28.031 | 47914 |
| 9 | Lysine | Malonylation | 86.000 | 50877 |
| 10 | Lysine | Succinylation | 100.016 | 49896 |
| 11 | Proline | Hydroxyproline | 15.995 | 46893 |
| 12 | Lysine | Glutarylation | 114.032 | 49197 |
| 13 | Lysine | GlyGlycylation | 114.043 | 47664 |
| 14 | Lysine | Hydroxyisobutylation | 86.037 | 49329 |
| 15 | Lysine | Propionylation | 56.026 | 49305 |
| 16 | Lysine | Trimetylation | 42.047 | 49296 |
| 17 | Arginine | Citrullination | 0.984 | 50087 |
| 18 | Arginine | Dimethylation_asym | 28.031 | 45183 |
| 19 | Arginine | Dimethylation_symm | 28.031 | 45666 |
| 20 | Arginine | Methylation | 14.016 | 45396 |
| 21 | Tyrosine | Nitrotyrosine | 44.985 | 48705 |

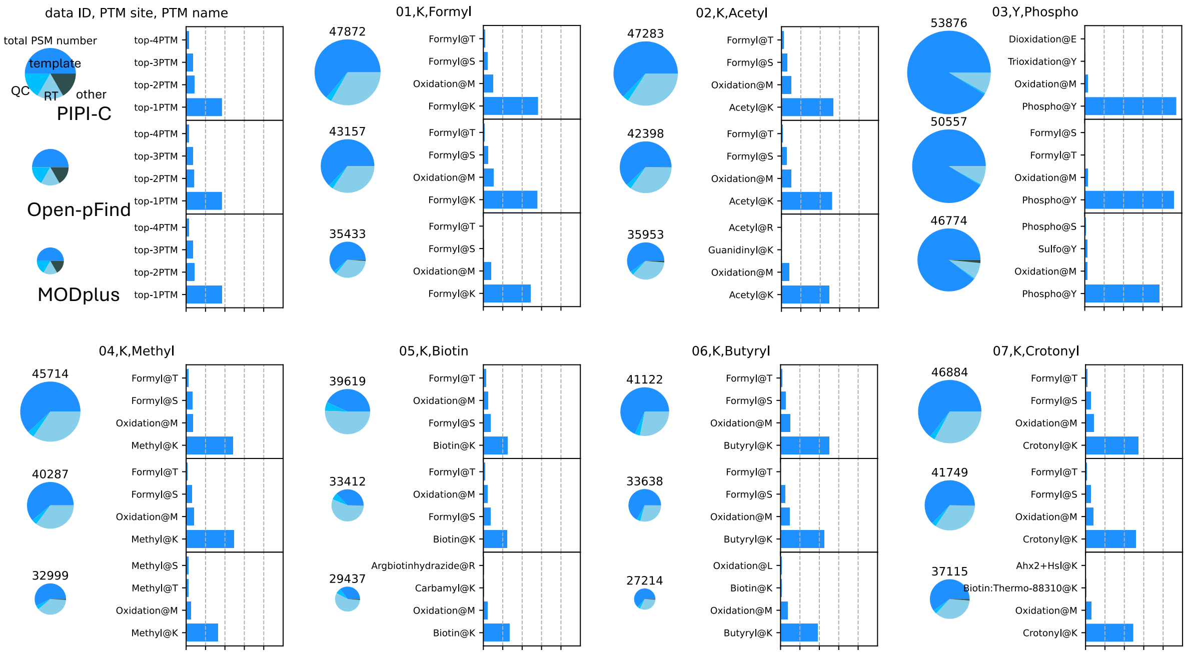


Figure S1: Comparison results of the synthetic data sets 01 to 07 at FDR of 0.01: bar chart, PSM numbers; pie chart, PTM numbers.

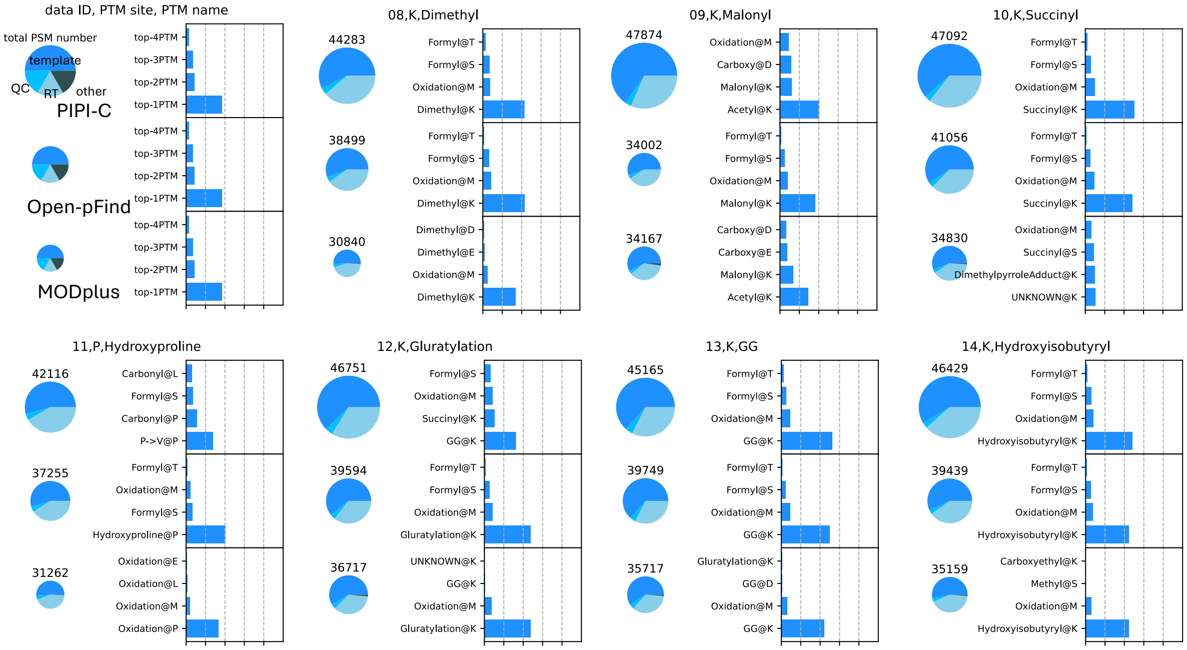


Figure S2: Comparison results of the synthetic data sets 08 to 14 at FDR of 0.01: bar chart, PSM numbers; pie chart, PTM numbers.

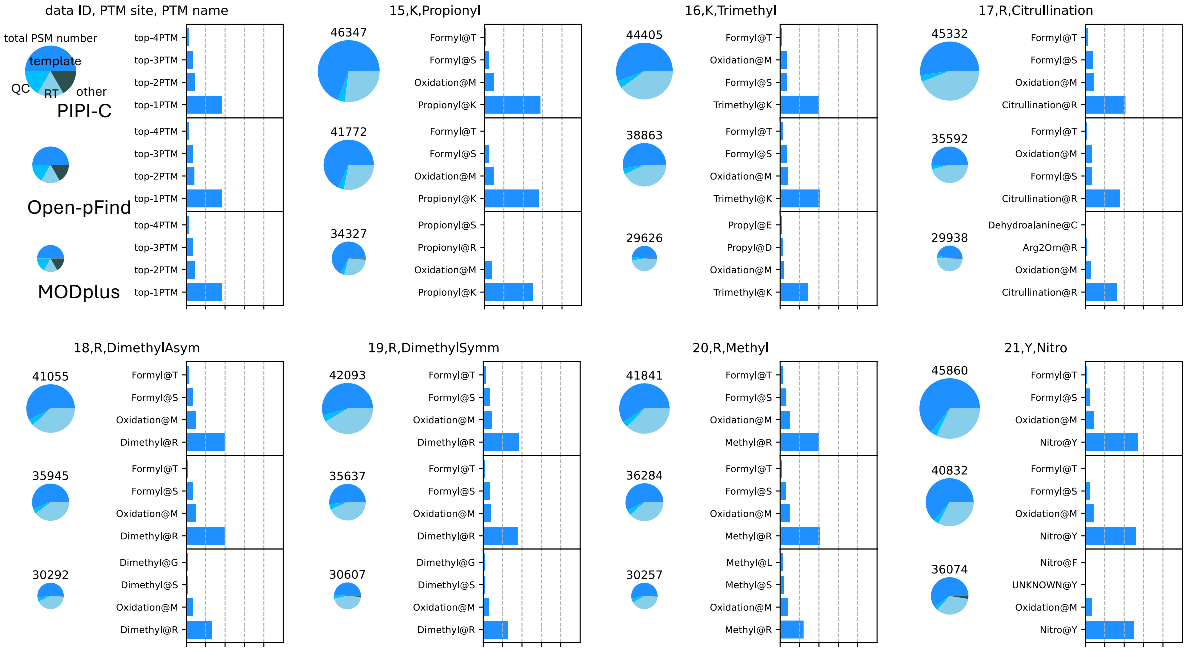


Figure S3: Comparison results of the synthetic data sets 15 to 21 at FDR of 0.01: bar chart, PSM numbers; pie chart, PTM numbers.

### 5 Supplementary Note 5: Results of replicated soybean data set

We repeated experiment 3 on five other replicated soybean data sets. The analysis is the same. As shown in **Figure** S4 to **Figure** S8, PIPI-C consistently outperform Open-pFind and MODplus in these replicate except R02.

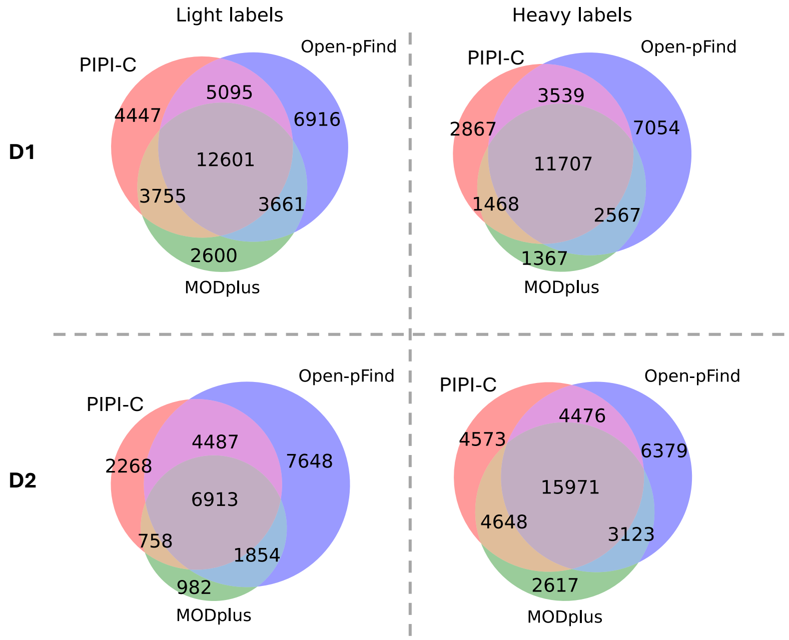


Figure S4: Replicate R02. Intersections of PSMs with fully dimethyl-labeled peptides identified from the soybean data sets. D1, D2: two samples in the data set.

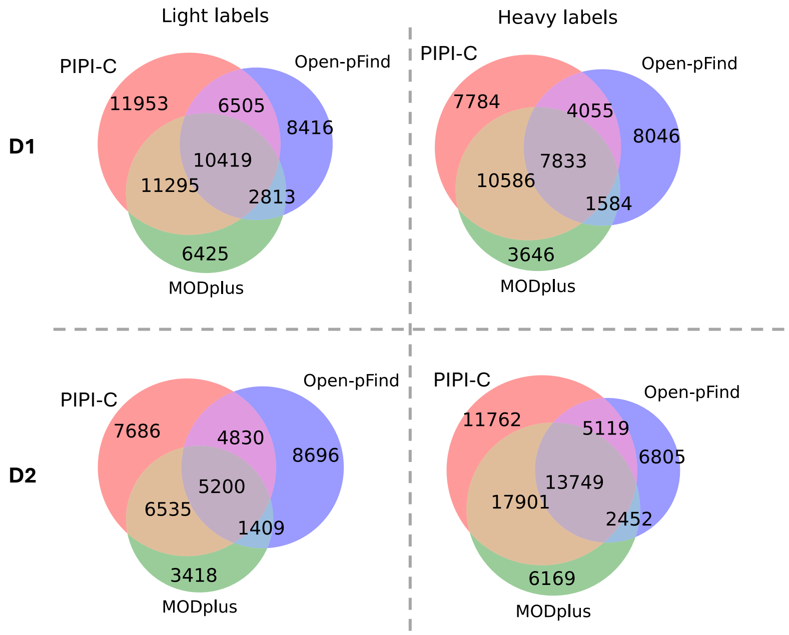


Figure S5: Replicate R03. Intersections of PSMs with fully dimethyl-labeled peptides identified from the soybean data sets. D1, D2: two samples in the data set.

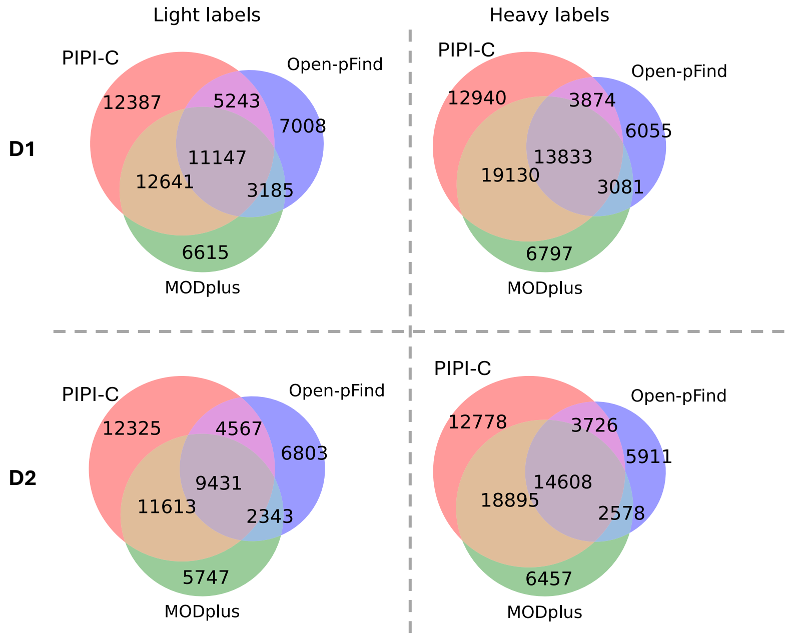


Figure S6: Replicate R04. Intersections of PSMs with fully dimethyl-labeled peptides identified from the soybean data sets. D1, D2: two samples in the data set.

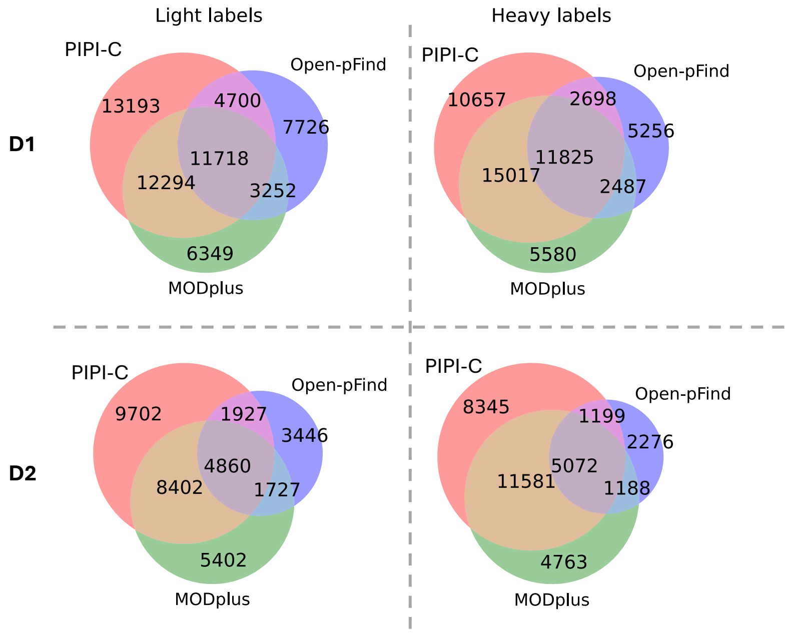


Figure S7: Replicate R05. Intersections of PSMs with fully dimethyl-labeled peptides identified from the soybean data sets. D1, D2: two samples in the data set.

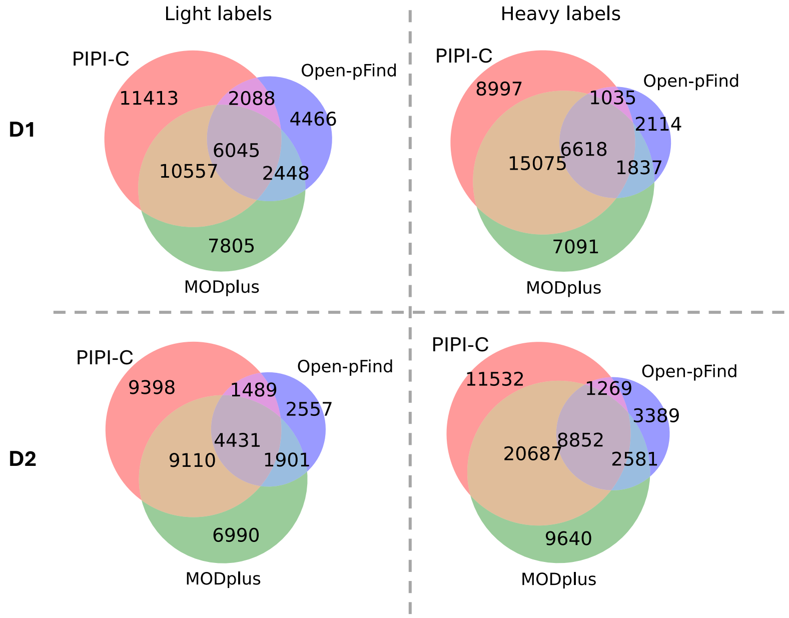


Figure S8: Replicate R06. Intersections of PSMs with fully dimethyl-labeled peptides identified from the soybean data sets. D1, D2: two samples in the data set.

### 6 Supplementary Note 6: More results of the *Petunia* data set

We recorded more results of the unidentified PTM combinations that can be identified when the PTM (other than GG@K) is pre-specified as a variable modification.

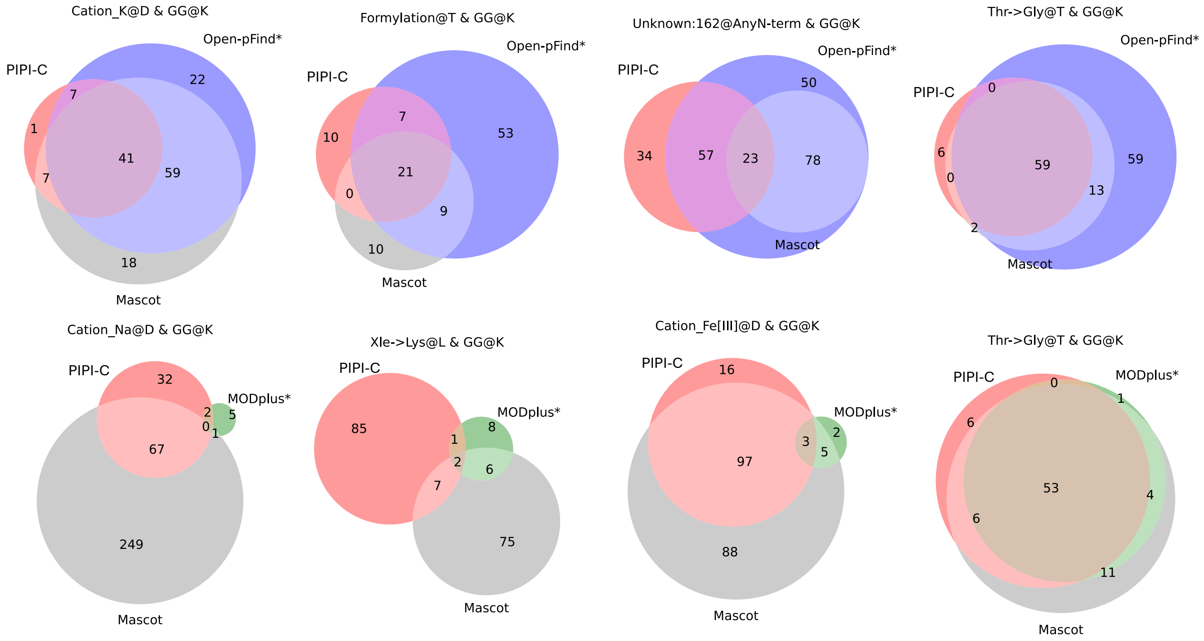


Figure S9: For Open-pFind and MODplus, unidentified PTM combinations can also be identified when the other PTM is pre-specified as variable modification
